# Designing stimuli for non-invasive, cell-type-specific manipulation of retinal ganglion cell activity

**DOI:** 10.64898/2026.09.30.755721

**Authors:** Niloufar Ghazavi, Saad Idrees, Vyom Manish Raval, Michael B. Manookin, Fred Rieke, Greg D. Field, Joel Zylberberg

## Abstract

The retina encodes visual information through parallel pathways formed by distinct retinal ganglion (RGC) types. Current understanding of how each cell type contributes to downstream visual processing and perception is limited by a lack of tools for selectively manipulating the activities of individual RGC types. Here, we introduce a computational framework called STORM (SpatioTemporal Orthogonal Response Modulation) that creates visual stimuli that aim to selectively modulate the activities of specific RGC types while minimally affecting the activities of nearby RGCs of non-targeted types. To illustrate the approach, demonstrate its flexibility, and identify conditions under which each simulated RGC type could be selectively manipulated, we first applied this framework to simplified models of RGC function, such as Linear-Nonlinear-Poisson (LNP) and LN-LN cascade models. We also deployed this approach on more complex (and more accurate) models of RGC responses, based on convolutional neural networks. We demonstrate that differences in spatial receptive fields, temporal dynamics, response polarities, and nonlinearities can provide sufficient degrees of freedom for designing stimuli that selectively manipulate individual RGC types. We also generalize this approach to conditions where the receptive field locations are unknown for in vivo and clinical applications.

## Introduction

Selective manipulation of individual cells and cell types is a major goal in neuroscience. This goal is motivated by the fact that the complexity of neural circuits requires causal manipulations to understand the relationships between neural activity, circuit architecture, perception, behavior, and disease [Panzeri et al., 2017, Fakhar et al., 2023]. The recognized importance of cell-type specific manipulations has led to many optogenetic and chemogenetic tools for activating or suppressing neuronal activity [Boyden et al., 2005, Fenno et al., 2011, Armbruster et al., 2007, Sternson and Roth, 2014]. However, these approaches frequently require the direct application of light or designer drugs to the brain. They also frequently require genetically modified organisms, such as mouse lines that express cre in a targeted cell type. These requirements limit the utility of these approaches in many experimental systems, particularly in non-human primate models and in human clinical applications.

Here we introduce STORM (SpatioTemporal Orthogonal Response Modulation), a method for cell-type specific manipulation of retinal neurons using designer visual stimuli. The retina has many well defined neuronal cell types [Wassle and Boycott, 1991, Field and Chichilnisky, 2007, Goetz et al., 2022] and the recent development of highly accurate models of retinal function facilitate these efforts [Maheswaranathan et al., 2023, Idrees et al., 2024, D’Agostino et al., 2025, McIntosh et al., 2016]. Many previous efforts have attempted to selectively manipulate the activity of visual neurons using light stimuli [Pokorny and Smith, 1997, Livingstone and Hubel, 1988]. A classic example is silent substitution, which uses the known spectral sensitivities of cone photoreceptors to design stimuli that selectively manipulate the activity of one or more cell types while being ’silent’ or ’metameric’ to the other types [Ishihara, 1906, Estévez and Spekreijse, 1982, Nugent et al., 2023]. More recently, large neural network models that accurately predict visually-evoked neural activity have been used to design stimuli that selectively manipulate individual neurons in the retina and in higher visual areas [Bashivan et al., 2019, Gaziv et al., 2025], and to identify stimuli that either maximally excite [Walker et al., 2019, Höfling et al., 2024, Burg et al., 2024], or parametrically drive [Prince et al., 2026] individual neurons or recording sites. However, these approaches did not target entire populations of specific cell types while holding fixed the responses of non-targeted cell types. Additionally, selective suppression has been less extensively tested than response enhancement. Finally, most previous studies used static natural images instead of movies. Here, we introduce an approach that overcomes these limitations.

To manipulate the activity of individual retinal cell types, we use models of stimulus-evoked neural responses to identify directions in stimulus space that change the activity of one or more cell types while minimally affecting non-targeted cell types. This allows the generation of stimulus pairs, in which the responses of one (or more) targeted cell types is modulated between presentation of stimulus pairs, while the responses of non targeted cell types are minimally affected (i.e., the stimuli are metameric to the non-targeted cell types). We applied this approach to diverse types of primate retinal ganglion cells (RGCs). To build intuition and demonstrate feasibility, we apply the approach to a series of models of increasing complexity, including: (1) Linear-Nonlinear-Poisson (LNP) models, (2) LN-LN cascade models, and (3) multilayer convolutional neural network (CNN) models. In simulations, we presented pairs of stimuli that were designed to change the activity of the targeted cell types, while remaining metameric to the non-targeted types. We used both natural and artificial stimuli as the starting point for generating these stimulus pairs. We show that selective manipulation is feasible for both naturalistic and artificial visual stimuli. STORM can exploit differences in both linear and nonlinear mechanisms to generate these stimulus pairs; selective manipulation is, in fact, easier to achieve in the presence of nonlinearities. Additionally, we extend this approach to achieve translation invariance, so that the approach is effective even without knowledge of RGC receptive field locations. Finally, we identify a range of conditions under which STORM is reliable. This study demonstrates that cell-type-specific manipulation of RGC activity is achievable with potential applications to studies of visual processing and disease diagnosis.

## Results

The intuition behind the approach for selectively manipulating the activities of specific RGC types is illustrated using a simplified example with two neurons and a two-pixel stimulus (Fig. 1A). The two neurons have different sensitivities to the two pixel intensities. This is summarized by the vectors representing the response gradient of each neuron with respect to changes in the pixel intensities. Changing the pixel values along these vectors increases the firing rate of the corresponding neuron, while moving in the opposite direction decreases the firing rate. The goal is to reduce the activity of one neuron (RGC A) without impacting the activity of the other (RGC B). To reduce the activity of RGC A, the stimulus is moved in the direction opposite to the gradient vector of RGC A in pixel space. Because this direction has a non-zero projection onto the gradient of RGC B, changing the pixel values in this direction will also affect the activity of RGC B. However, by removing the component of the pixel modification that affects RGC B, we obtain directions that modulate RGC A without changing the response of RGC B (Fig. 1A, magenta and blue arrows).

**Figure 1.**
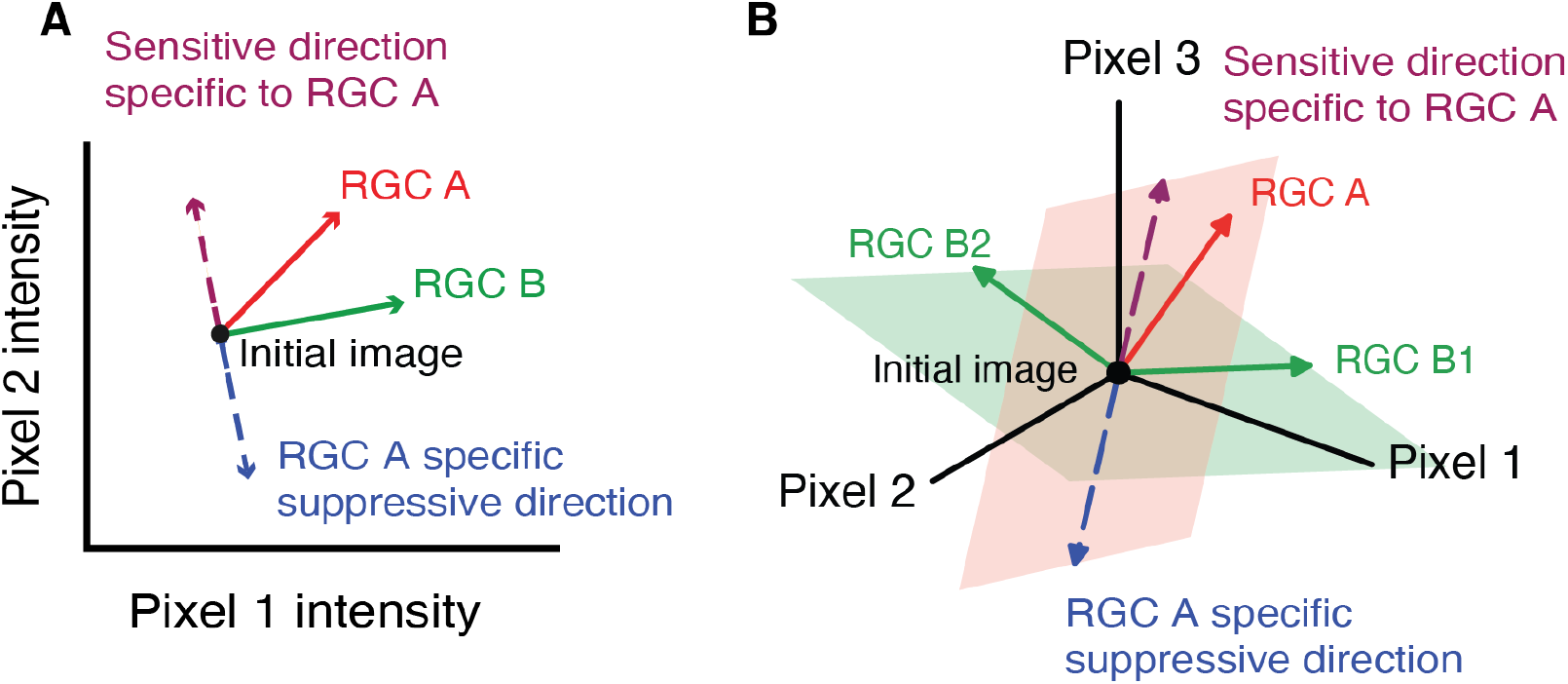
SpatioTemporal Orthogonal Response Modulation (STORM) framework for selective manipulation of RGC activities. **(A)** Illustration for two RGCs (A and B) driven by a two-pixel stimulus. The black dot represents the initial (unmodified) stimulus. Solid arrows represent the direction of maximum sensitivity or ’firing rate gradients’ of RGC A (red) and RGC B (green) with respect to the pixel values. Dashed magenta and blue vectors indicate the directions along which moving the stimulus selectively increases or decreases the response of RGC A, without affecting the response of RGC B. **(B)** Generalization of the approach to higher-dimensional stimulus spaces and multiple non-target RGCs. The green hyperplane represents the subspace spanned by the gradients of the non-target RGCs B1 and B2. The red hyperplane illustrates the subspace containing the sensitive direction of the target RGC A. The magenta and blue vectors indicate directions along which the stimulus can be moved to selectively modulate the response of RGC A while leaving the responses of RGCs B1 and B2 unaffected.

This approach, which we call STORM, generalizes to more complex geometries and conditions. For example, in Fig. 1B, RGCs B1 and B2 represent neurons of one type and RGC A represents a neuron of a different type. To find a stimulus perturbation that will decrease the activity of RGC A without changing the activities of B1 and B2, we use the response gradients of non-target RGCs (B1 and B2) to generate an orthonormal basis, a set of perpendicular, unit-length directions that spans the stimulus directions that affect the responses of these RGCs (see Methods). By projecting the gradient of RGC A onto this orthonormal basis and subtracting this projection from its original gradient, one can identify a direction in stimulus space that selectively manipulates RGC A while being metameric to RGCs B1 and B2 (Fig. 1B). Such selective manipulation will always be possible if the gradients for RGCs of different types are linearly independent in the stimulus space. Furthermore, the stimulus space must be sufficiently high-dimensional to identify directions that selectively manipulate a target neuron while leaving non-target responses unchanged.

The remainder of this paper tests the generality of STORM by applying it to multiple cell types, a larger stimulus space, and neurons with increasingly nonlinear stimulus-response relationships to show the generality of the approach. We focus on the four numerically dominant output neurons of the primate retina: ON and OFF parasol and ON and OFF midget RGCs. These four types constitute ~70% of the output of the primate retina [Dacey, 2004]. They are irreducible cell types: each tiles the retina in a mosaic-like pattern to fully sample visual space [Wassle and Boycott, 1991, Devries and Baylor, 1997, Field and Chichilnisky, 2007, Roy et al., 2021]. This means that each visual field location is sampled by all four types. Furthermore, their spatiotemporal receptive fields overlap considerably. Thus, these four RGC types provide a challenging and well-defined case for testing the proposed framework for generating visual stimuli that specifically manipulate targeted cell-types.

### Cell-type specific manipulations are achievable with linear-nonlinear model neurons

To examine how well the approach outlined in Fig. 1 could selectively manipulate RGC responses, we constructed four simulated populations of ON and OFF parasol and midget RGCs using linear-nonlinear Poisson (LNP) models (Fig. 2A). Model parameters were chosen to mimic ON and OFF parasol and midget RGCs [Parthasarathy et al., 2017, Kling et al., 2020]. For example, ON and OFF parasol RGCs were modeled to have larger receptive fields with briefer and more biphasic temporal filters than ON and OFF midget RGCs (Supplementary Fig. S1B).

**Figure 2.**
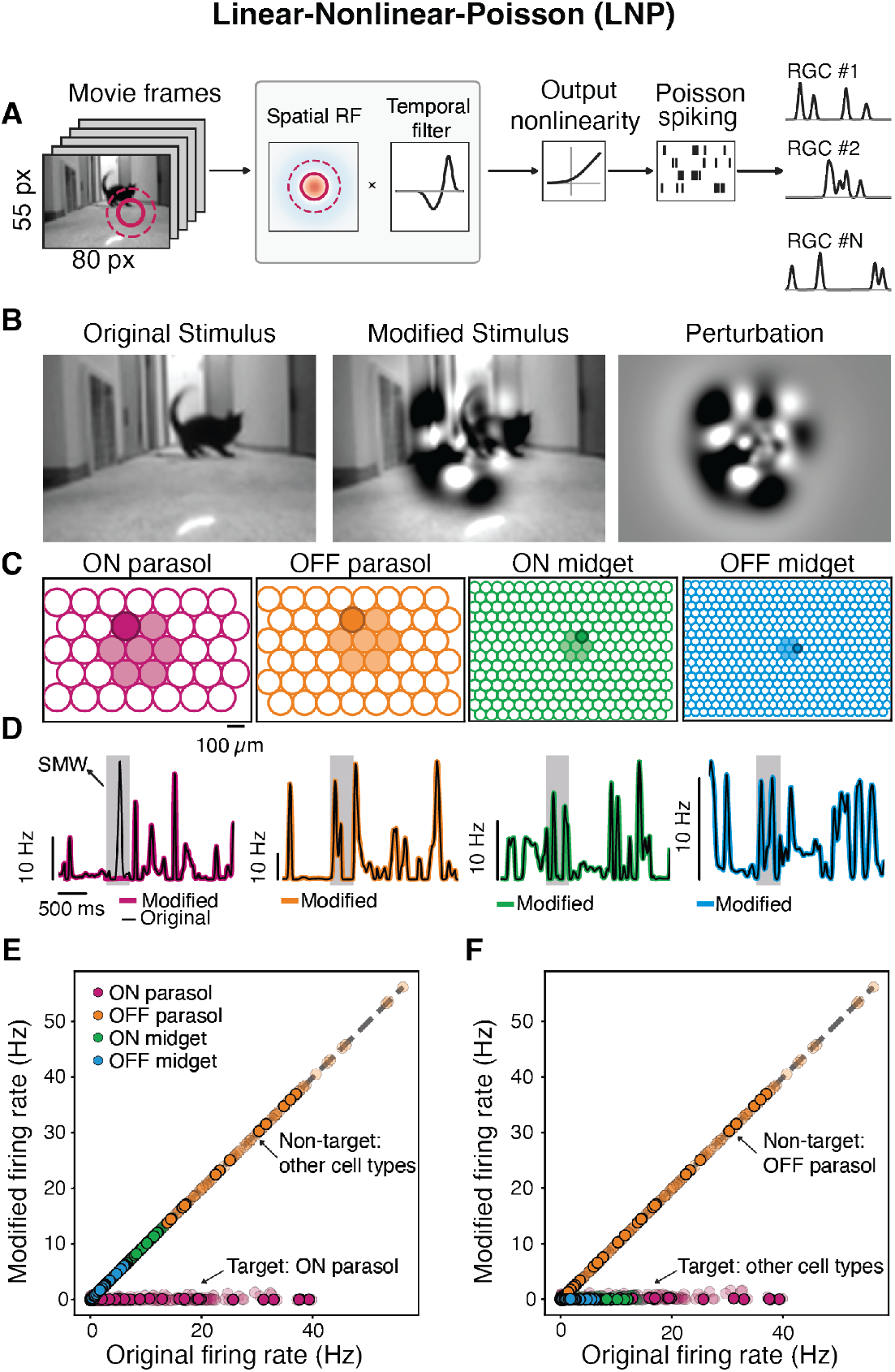
Selective manipulation of simulated RGC responses using a Linear–Nonlinear–Poisson model. **(A)** Schematic of the Linear–Nonlinear–Poisson (LNP) model. A visual stimulus is filtered by spatiotemporal receptive field (RF). The resulting generator signal is converted to an instantaneous firing rate by a static nonlinearity and converted to discrete spike counts by a Poisson process. **(B)** Example frame of the original natural movie (left), the corresponding frame of the modified movie that suppresses ON parasol cells (middle), and the difference between the modified and original frames (perturbation; right). **(C)** RF mosaics of the simulated RGC types, each RF drawn at its 1-*σ* center contour. Filled RFs indicate the cells included in the manipulation; darker cells are shown in **D. (D)** Firing rates for example cells in response to the original (black) and modified (colored) movies; the stimulus manipulation window (SMW) is shaded gray. The ON parasol response is reduced during the SMW; the non-target responses are unmodified. **(E)** Firing rate of every cell at each time bin (each point) within the SMW for the original vs modified movies. ON parasol cells are suppressed to near-zero rates; the three non-target types are unmodulated. Opaque points are the four cells shown in **D. (F)** Same as **E**, but for simultaneous suppression of three RGC types: ON parasol, ON midget and OFF midget. OFF parasol RGCs are the non-targeted type. All targeted RGCs are driven to near-zero firing within the SMW; OFF parasol responses are unmodified. Corresponding stimulus frames and firing rates are shown in Supplementary Fig. S3 **B, D**.

We simulated responses of these neurons to a naturalistic movie (Fig. 2B) and computed the response gradients of all neurons with shaded receptive fields with respect to each pixel value in the movie. Note that the receptive fields of the targeted and non-targeted cell types overlapped (Fig. 2C). In this first example, we selectively manipulated the activities of ON parasol cells. The gradients were used to construct an orthonormal basis spanning stimulus directions affecting the non-target cell types (ON midget, OFF midget and OFF parasol). Removing the component of the ON parasol response gradient that aligns with this basis yielded a direction in stimulus space that selectively manipulates the activities of the ON parasol RGCs. As expected, moving opposite to this direction selectively reduced the activity of the simulated ON parasol RGCs while remaining metameric to the other three RGC types, such that their responses were minimally changed relative to the original stimulus (Fig. 2D,E) (see Methods). In fact, perturbing the stimulus sufficiently in this direction yielded a near-complete suppression of the simulated ON parasol cell responses with minimal impact on the firing rates of the non-target cell types (Fig. 2E).

We similarly identified a direction in stimulus space that selectively suppressed ON midget responses while keeping the modified stimulus metameric for the other three RGC types (Supplementary Fig. S2). We also tested the extent to which the responses of three simulated RGC types (ON parasol and ON and OFF midget) could be selectively suppressed while remaining metameric for the fourth (OFF parasol). This was likewise achievable (Fig. 2F; Supplementary Fig. S3).

### Cell-type specific manipulations are achievable with more complex neural models

LNP models fail to capture a variety of nonlinearities in RGC responses [Enroth-Cugell and Robson, 1966, Schwartz et al., 2012, Freeman et al., 2015]. Do such nonlinearities limit the ability of STORM to generate stimuli that selectively manipulate the activity of specific cell types? To answer this question, we first deployed a cascaded LN-LN model (Fig. 3A) that is commonly used for capturing a key set of so-called ‘subunit’ nonlinearities in the retina [Hochstein and Shapley, 1976, Gollisch and Meister, 2010, Meister, 2026]. Using this model, we again generated four simulated RGC types: consistent with the literature, the simulated ON and OFF parasol cells were assigned strong subunit rectification, while the simulated ON and OFF midget cells had weaker subunit rectification [Petrusca et al., 2007, Cafaro and Rieke, 2013]. We then repeated the analysis performed using the LNP models to find a direction in stimulus space that would suppress the simulated ON parasol RGCs while being metameric to the other three RGC types for these LN-LN models (Fig. 3B). We again focused on the central cluster of cells with the shaded receptive fields (Fig. 3C).

**Figure 3.**
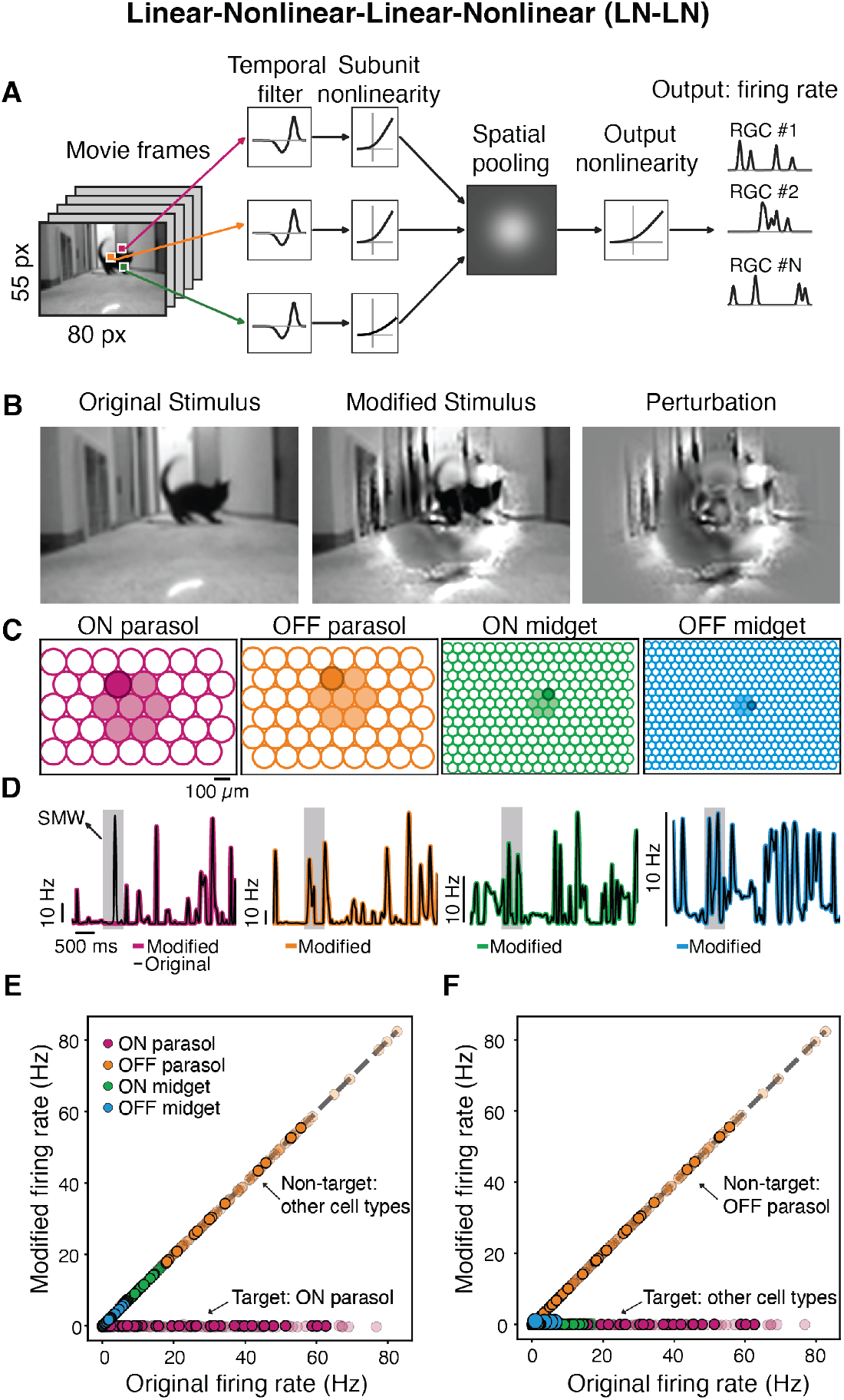
Selective manipulation of simulated RGC responses using a LN-LN model. **(A)** Schematic of the LN-LN model. The movie is filtered in time at every pixel and then passed through a subunit nonlinearity applied to each pixel. Subunit outputs are pooled over space with a Gaussian RF, and passed through an output nonlinearity to yield instantaneous firing rates. **(B)** Example frame of the original natural movie (left), the corresponding frame of the modified movie that suppresses ON parasol cells (middle), and the difference (perturbation, right). **(C)** RF mosaics of the simulated RGC types drawn at 1-*σ*. Filled RFs indicate the cells included in the manipulation; darker cells are shown in **D.(D)** Firing rates for example cells in response to the original (black) and modified (colored) movies; the stimulus manipulation window (SMW) is shaded gray. The ON parasol response is reduced during the SMW; the non-target responses are unmodified. **(E)** Firing rate of every cell at each time bin (each point) within the SMW for the original vs modified movies. ON parasol cells are suppressed to near-zero rates; the three non-target types are unmodulated. Opaque points are the four cells shown in **D. (F)** Same as **E**, but for simultaneous suppression of three RGC types: ON parasol, ON midget and OFF midget. OFF parasol RGCs are the non-targeted type. All targeted RGCs are driven to near-zero firing within the SMW; OFF parasol responses are unmodified. Corresponding stimulus frames and firing rates are shown in Supplementary Fig. S4 **B, D**.

In contrast to the LNP model (Fig. 2) where the gradient direction of each RGC is fixed by its linear receptive field, in the LN-LN model, the gradient direction depends on the stimulus. This is because the nonlinear subunits weight different regions of the receptive field differently depending on the stimulus, causing the gradient direction to change as the stimulus is modified. Thus, we used an iterative approach wherein the stimulus was modified incrementally, and the gradient directions were re-calculated at each iteration (see Methods). This iterative procedure identified a stimulus modification that silenced the ON parasol cells while being metameric for the other three RGC types (Fig. 3B, D, E). Selective suppression of three RGC types was also achieved with the LN-LN model (Fig. 3F; Supplementary Fig. S4).

Subunit nonlinearities are just one of several nonlinear mechanisms found in the retina [Baccus and Meister, 2002, Berry et al., 1999, Gollisch and Meister, 2010, Kim and Rieke, 2001]. For a more general test of the approach and its robustness to nonlinearities, we fit a deep convolutional neural network (CNN) model (Fig. 4A) to the measured responses of ON and OFF parasol and midget cells to white noise checkerboard and natural scenes with superimposed eye movements, recorded simultaneously with large-scale multi-electrode arrays (MEAs) [Idrees et al., 2024] (see Methods). The CNN well approximated the responses of all four RGC types (Supplementary Fig. S5).

**Figure 4.**
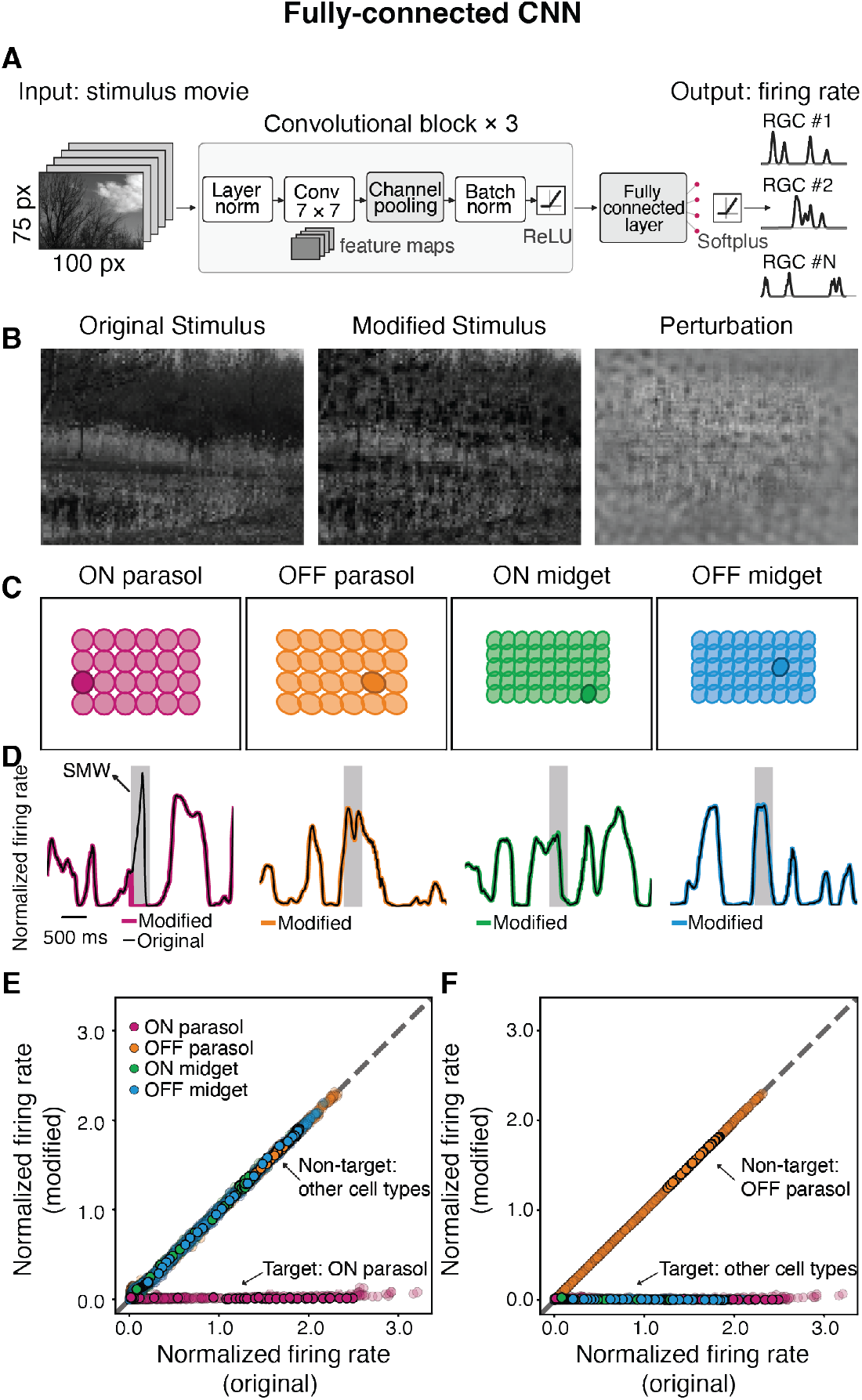
Selective suppression of simulated RGC responses using a CNN model. **(A)** Schematic of the CNN model. The movie is passed through a series of convolutional layers that extract visual features, followed by a fully connected layer and output nonlinearity that predict the instantaneous firing rate of each RGC. **(B)** Example frame of the original natural movie (left), the corresponding frame of the modified movie that suppresses ON parasol cells (middle), and the difference (perturbation, right). **(C)** RF mosaics of the simulated types drawn at 1-*σ*. Filled RFs indicate the cells included in the manipulation; darker cells are shown in **D.(D)**Firing rates normalized by the median firing rate for example cells in response to the original (black) and modified (colored) movies; the stimulus manipulation window (SMW) is shaded gray. The ON parasol response is reduced during the SMW; the non-target responses are unmodified. **(E)** Firing rate of every cell at each time bin (each point) within the SMW for the original vs modified movies. ON parasol cells are suppressed to near-zero rates; the three non-target types are unmodulated. Opaque points are the four cells shown in **D. (F)** Same as **E**, but for simultaneous suppression of three RGC types: ON parasol, ON midget and OFF midget. OFF parasol RGCs are the non-targeted type. All targeted RGCs are driven to near-zero firing within the SMW; OFF parasol responses are unmodified. Corresponding stimulus frames and firing rates are shown in Supplementary Fig. S6 **D**,**E**.

Given that this CNN model exhibited comparable performance to other recent models of the primate retina [Idrees et al., 2024, Gogliettino et al., 2024, Shah et al., 2022], we used it to investigate the ability of STORM to find stimulus pairs that selectively change the activity of one or more cell types to a natural movie stimulus (Fig. 4B). We constructed homogeneous mosaics of each RGC type that tiled space using the CNN (Fig. 4C)(see Methods) and then used the gradients of the responses with respect to pixel intensities to find a stimulus modification that would suppress the activity of the ON parasol cells while being metameric to the other three RGC types. Using this procedure, we were again able to find a stimulus perturbation (Fig. 4B) that would suppress the population of ON parasol cells (Fig. 4D, E). Three RGC types could likewise be suppressed simultaneously with the CNN model (Fig. 4F; Supplementary Fig. S6D, E). Selective suppression of ON parasol cells was also achieved when the initial stimulus was a white-noise checkerboard (Supplementary Fig. S7)

In the simulations presented thus far, cells of a given RGC type were identical except for their receptive-field locations. However, cells of the same type can exhibit variability in their response properties [Gauthier et al., 2009, Shah et al., 2022]. We therefore sought to determine the extent to which the approach remained effective in the presence of such within-type heterogeneity. To answer this question, we repeated the analysis using CNN models that included within-type heterogeneity from measured RGC responses (Supplementary Fig. S8 C) (see Methods). The cell-to-cell variability did not diminish our ability to selectively suppress the activity of targeted RGC populations while remaining metameric to the other types (Supplementary Figs. S8, S9).

In summary, nonlinearities and cellular heterogeneity do not diminish the ability of STORM to find stimulus pairs that manipulate the activity of one or more RGC types while being metameric for non-targeted RGC types. We provide an intuition for this result and identify potential failure modes in the Discussion. Next, we show that cell-type-specific manipulation of RGC activity remains achievable even without knowledge of individual RGC receptive field locations.

### Cell-type specific manipulations are achievable with translation-invariant models

The preceding demonstrations that stimulus modifications can selectively modulate the firing rates of specific RGCs relied on idealized conditions in which the spatial locations of individual RGC receptive fields were known precisely (Figs. 2, 3, 4). However, in many practical applications, this information will not be available like when presenting these stimuli to awake, behaving animals or to human subjects.

This limitation led us to ask whether RGC type-specific manipulation could be achieved without knowledge of individual receptive-field locations. To answer that question, we developed a translation-invariant CNN model of RGCs (Fig. 5A). Unlike the models which predicted the firing rates of individual RGCs, this model outputs a set of firing rate maps—one for each RGC type. Just as the input is a movie, each firing-rate map is also a movie with matched spatiotemporal dimensions (Fig. 5B). Each map is generated using a shared set of RGC-type-specific weights, trained jointly on data from all RGCs of that type across the visual field, rather than fit independently to each cell. These shared weights are convolutionally applied across space. Because the weights are reused at every location, the model effectively oversamples the visual field by evaluating the response at each position in the map. This makes the model inherently translation-invariant.

**Figure 5.**
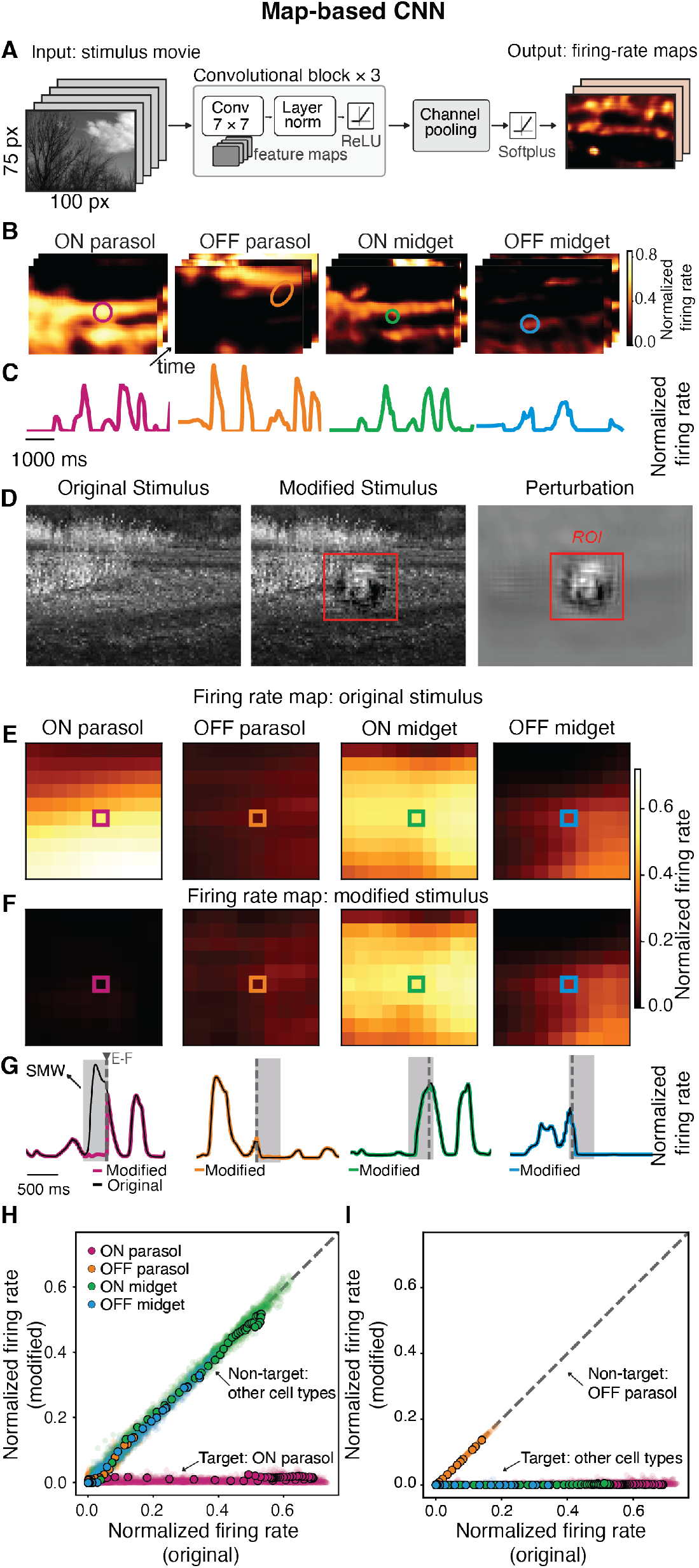
Selective RGC manipulation with translation invariance using a CNN and firing rate maps. **(A)** Schematic of the map-based CNN. The convolutional framework matches Fig. 4 but without a fully-connected readout. The output is a firing-rate map of same spatial dimensions as the input movie, with one channel per RGC type. Thus, the model predicts the response of a cell of a given type at any location, rather than only specific locations. **(B)** Example firing-rate maps for the four RGC types at successive time points. The colored circles mark RF centers over which a firing rate can be integrated. **(C)** Predicted firing rate over time at the RF locations in **B. (D)**Example frame of the original natural movie (left), the corresponding frame of the modified movie that suppresses ON parasol cells (middle), and the difference (perturbation, right). The outlined region denotes the region of interest (ROI) analyzed in **E-I.(E)**Firing-rate maps within the ROIs of **D** for each RGC type in response to the original movie, at the time indicated by the dashed vertical line in **G. (F)** Same as **E**, for the modified movie. The ON parasol firing rate map is driven to near-zero within the ROI, whereas the other three non-target RGC types are unaffected. **(G)** Predicted firing rate over time at the locations marked by the colored squares in **E-F**, normalized by the maximum firing rate, in response to the original (black) and modified (colored) movies. Gray shading marks the SMW; the dashed vertical line marks the time point of the maps shown in **E-F. (H)** Scatter plot compares firing rate during the SMW at every location within the ROI, in response to the original and the modified movie. ON parasol responses are suppressed at all locations; the non-target RGC responses are unchanged. Opaque points are the four cells shown in **G. (I)** Same as **H** for simultaneous suppression of ON parasol, ON midget and OFF midget cells; OFF parasol is the non-target cell type. The corresponding stimulus frames and firing rates are shown in Supplementary Fig. S12 B-F.

This mapped-based model was fit to predict RGC responses to checkerboard noise and natural scenes with superimposed eye movements (see Methods). Fit quality was high and on par with other recently developed fitting approaches (Supplementary Fig. S10) [Idrees et al., 2024].

We generated stimulus pairs that manipulated targeted RGC types, while being metameric to non-targeted types (Fig. 5D). The stimulus modifications are formulated at the level of the maps rather than individual RGCs. To selectively suppress ON parasol RGC responses, we designated a 10×10 pixel region of interest within each firing-rate map, yielding 100 sampled spatial locations (Fig. 5D-F). For each of these locations, we computed the gradient of the predicted firing rate with respect to every pixel of the input movie. We then applied the same procedure used for the previous models to identify an RGC-type-specific sensitive direction in stimulus space. Moving opposite to this direction selectively reduced the predicted map of ON parasol RGC responses (Fig. 5E-H), whereas moving along this direction enhanced their responses (Supplementary Fig. S11 C-F). The procedure also was able to generate stimulus pairs to manipulate the activity of three RGC types while remaining metameric to the fourth (Fig. 5I, Supplementary Fig. S12). In addition, suppression of one or several cell types were obtained when a white noise checkerboard was used as the initial stimulus instead of a natural movie (Supplementary Figs. S13, S14).These results demonstrate the potential to use this map-based approach in applications where receptive-field locations cannot be mapped or known in advance.

### Robustness to imperfect models

The STORM approach that we have described requires a model of RGC encoding to estimate the gradients for manipulating one RGC type and remaining metameric to the other types. How accurate must the model be for this procedure to succeed? To answer this, we generated a ‘tester’ model using the map-based CNN (Fig. 5) and generated an ensemble of ‘generator’ models that were spread along a continuum in terms of how accurately they could predict the stimulus-evoked responses of the ‘tester’ model (Fig. 6A). We created these ‘generator’ models by freezing model parameters at different stages of training (see Methods). Then, each ‘generator’ model was used to generate a stimulus pair that selectively suppressed the target RGC type until its mean firing rate reached 0.001 Hz. The modified stimulus from that generator model was then shown to the ‘tester’ model. If the stimulus from the (imperfect) generator model silenced the targeted RGC types in the tester model, that indicated that the generator model was sufficiently accurate (compared with the tester model) to reliably generate stimuli that selectivity manipulated the activity of targeted cell-types. The similarity between each generator model and the tester model was quantified as the model-to-model correlation of their predicted responses.

**Figure 6.**
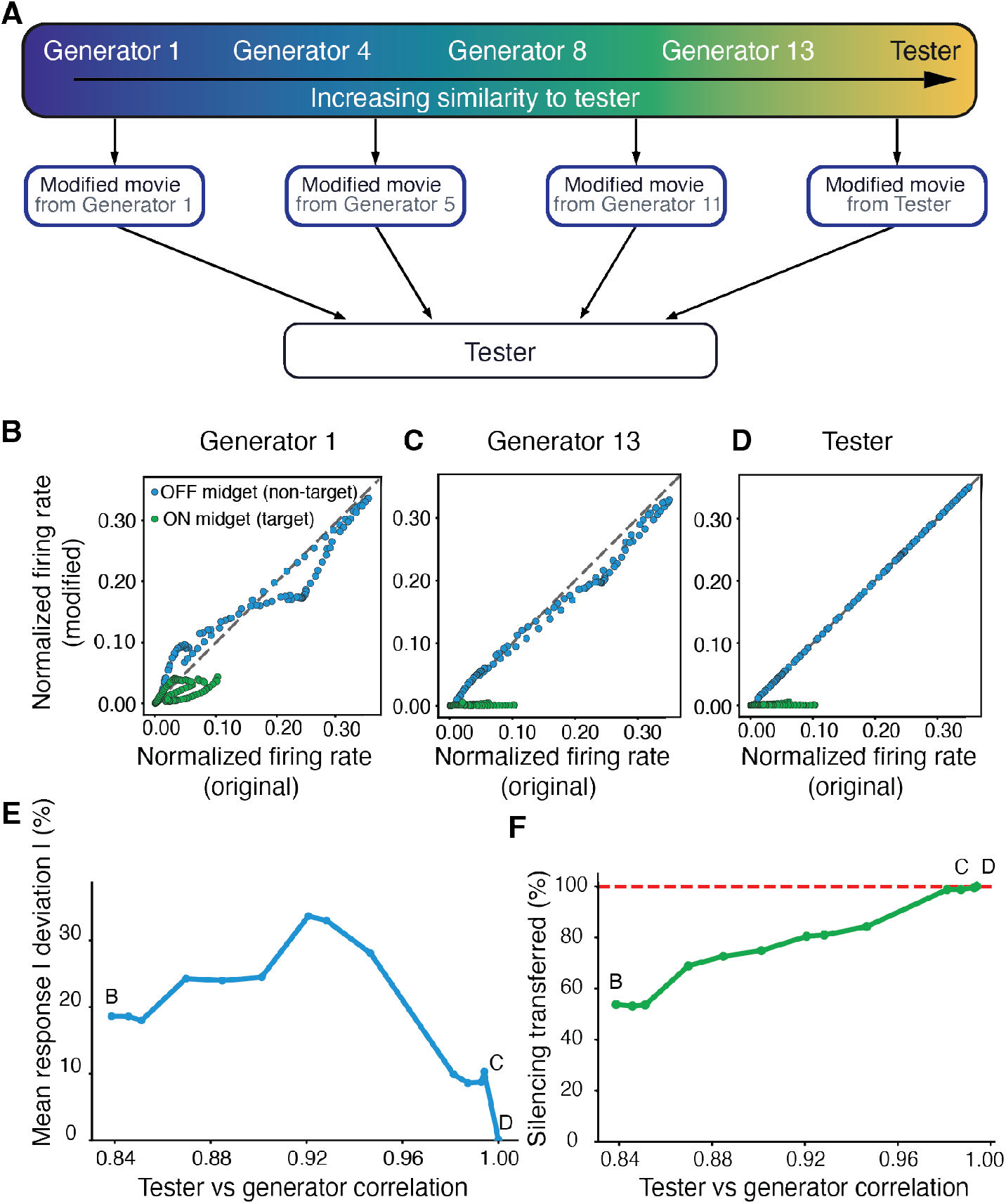
Robustness of selective manipulation to imperfect models. **(A)** Schematic of the robustness analysis. ‘Generator models’ with increasing predictive performance relative to the map-based CNN were each used to design a modified movie for suppressing ON midget RGCs without affecting OFF midget RGCs. Every modified movie was then evaluated against the single (best) ‘tester’ model. Similarity between each generator and the tester was quantified as the Pearson correlation between their predicted responses. **(B-D)** Predicted firing rates for the modified (y-axis) versus the original movie (x-axis), evaluated by the tester model, for ON midget (target, green) and OFF midget (non-target, blue) RGCs. The perturbation was designed by the generator least similar to the tester (**B**, Generator 1), by the generator most similar to the tester (**C**, Generator 13), and by the tester model itself (textbfD). **(E)** *Stimulus selectivity*, quantified as the mean absolute percentage change in non-target firing rate relative to the original rate, plotted against the correlation between generator and tester predictions. Letters mark the generators shown in **B–D**. Non-target deviations were substantial for generators that weakly correlate with the tester and fall toward zero as the correlation approaches 1. **(F)** *Stimulus efficacy*, quantified as the reduction in target firing rate achieved using the tester model, normalized by the corresponding reduction in the generator model. The percentage of target suppression transfer increases as the generator becomes more similar to the tester.

With relatively dissimilar generator models (correlation ~84–85%), targeted cell responses (ON midget RGCs) were suppressed by approximately 50–55%, while non-target responses (OFF midget) altered by 18–20% on average (Fig. 6B,E,F). As generator– tester similarity increased, suppression of the responses of the targeted cells increased, reaching approximately 80% at a correlation of 92% and nearly complete suppression at correlations above 98% (Fig. 6F). For the non-target cells, responses progressively approached the unity line at high generator–tester similarity (Fig. 6C). The mean absolute deviation of non-target responses was non-monotonic, it increased to about 33% near a correlation of 92% before decreasing to about 10% at correlations above 98% (Fig. 6E). When the tester itself was used to generate the modified movie, the target cell responses were completely suppressed, while the responses of the non-target cells were unaffected. These results indicate the model accuracy needs to be reasonably high, but not perfect, to strongly modulate the activity of the targeted cell type, while remaining metameric to non-targeted types.

## Discussion

In this study, we established STORM (SpatioTemporal Orthogonal Response Modulation), a computational framework for designing visual stimuli that selectively and noninvasively manipulate the activity of specific retinal ganglion cell (RGC) types while minimizing the effects on non-target cell types. Using synthetic RGC populations, we found that gradient-based stimulus optimization can identify perturbations that selectively manipulate the responses of target RGC types. A central conceptual finding is that differences in spatial receptive fields, temporal dynamics, nonlinearities, and response polarity across RGC types provide sufficient degrees of freedom to enable selective control.

The proposed framework generalizes the classic silent substitution technique [Estévez and Spekreijse, 1982, Nugent et al., 2023], which isolates specific photoreceptor pathways (e.g., L-cones vs M-cones) by identifying light mixtures that are orthogonal to the spectral sensitivities of non-target cells [Derrington et al., 1984]. We extended this principle from the spectral domain to the high-dimensional spatiotemporal domain, demonstrating that analogous orthogonal stimulus subspaces can be used to selectively suppress or enhance the responses of particular RGC types.

Deep learning has led to the generation of increasingly accurate models of neural response to visual stimuli [Yamins et al., 2014, McIntosh et al., 2016, Idrees et al., 2024, Gogliettino et al., 2024]. These models have been used in several contexts to design more effective stimuli for driving or differentiating between neural responses. For example, ’Most Exciting Input’ (MEI) [Walker et al., 2019, Höfling et al., 2024, Pierzchlewicz et al., 2023] and Extending DeepDream with Real-time Evolution for Activity Maximization (XDREAM) [Ponce et al., 2019], synthesize stimuli that maximize the response of a single cell or cell type, but impose no explicit constraint on the responses of other cells or cell types. ‘Most Discriminative Stimuli’ (MDS) is designed for cell-type-based activation but is not designed to generate stimulus pairs that are metameric for non-target cell types [Burg et al., 2024]. By guaranteeing that non-target types remain unaffected, the orthogonalization procedure we present makes it possible to attribute any downstream effect specifically to the manipulated population. Accurate neural encoding models have also been used to design natural image perturbations that selectively modulate activity in a targeted recording site along arbitrary, experimenter-chosen directions while minimizing changes in non-target recording sites [Gaziv et al., 2025]. However, in that approach, complete invariance of responses at non-target recording sites was not achieved and neural silencing was challenging. By contrast, STORM enables manipulation of all RGCs within a targeted cell type and this could be achieved through response suppression or enhancement. Finally, previous model-guided stimulus design studies have synthesized stimuli *de novo*, without reference to any particular natural scene, focusing predominantly on generating static stimuli [Walker et al., 2019, Bashivan et al., 2019, Ponce et al., 2019], with recent extensions to artificial spatio-temporal stimuli [Höfling et al., 2024],whereas STORM modifies natural movies themselves across both space and time. Thus, we view the direct orthogonalization approach presented here as complementary to other similar efforts, but it also presents several distinct advantages in potentially minimizing off-target effects, generating more complex stimuli, and improving interoperability.

Importantly, we found that STORM remains effective across a broad range of neural models, from simple synthetic populations to more realistic conditions with uncertainty in receptive-field locations, including translations of the visual stimulus relative to the retinal mosaic. The framework was also robust to inaccuracies in the underlying neural response models and to cell-to-cell variability within RGC types. These results suggest that selective manipulation of retinal output through visual stimulation may remain feasible even under the imperfect conditions encountered in experimental and clinical settings.

### Nonlinearities provide additional leverage for selective manipulation

We showed that the inclusion of increasingly complex nonlinearities among RGC responses do not diminish the ability of the orthogonalization procedure to find stimuli that manipulate the activity of one or more cell types while being metameric for non-targeted cell types (Figs. 2, 3, 4). We develop some intuition for this potentially counterintuitive result.

For purely linear neurons, the response of each cell depends only on the inner product between its spatiotemporal receptive field (RF) and the stimulus. Thus, the linear receptive fields define both the stimulus directions that modulate a target neuron and the directions that must be avoided to keep the responses of non-target cells unchanged. In this case, selective stimulation is always possible when the receptive fields of the target population are not wholly contained within the linear span of the receptive fields of the non-target population.

Increasing model complexity to include a single output nonlinearity produces linear-nonlinear neurons (e.g., LNP neurons, Fig. 2). In this case, the gradient of the firing rate with respect to the stimulus is simply the linear receptive field, multiplied by a gain term that is equal to the slope of the nonlinearity (see Methods). Thus, the effective constraint on stimulus modification imposed by each neuron depends on the gain of its nonlinear response. Importantly, this means that introducing nonlinearities with rectification, such as ReLU or softplus functions, actually *simplifies* the selective manipulation of RGCs. When the generator signal (the similarity between the stimulus and the receptive field) is far below the nonlinearity’s threshold, its nonlinear gain vanishes and thus so does its gradient with respect to the stimulus. In this case, the neuron is insensitive to small changes in the stimulus and it effectively drops out of the set of constraints that need to be considered when designing RGC-specific stimulus modifications. Achieving selective RGC manipulation in the case of linear neurons requires satisfying the constraints imposed by *every* neuron, whereas achieving the same result in the LN case requires that only a subset of those constraints be satisfied. Consequently, whenever selective cell manipulation is achievable in the linear case, output nonlinearities can make such manipulation easier to achieve.

Including subunit nonlinearities, as in the LN-LN cascade model, further strengthens this effect because each subunit nonlinearity provides a new way to remove one of the constraints imposed on the stimulus manipulation. The CNN models, consisting of cascading sets of linear-nonlinear operations present an even more extreme version of this same idea. Thus, nonlinearities can, and frequently do, make selective RGC manipulation easier to achieve.

### Limitations and technical considerations of the gradient-based stimulus design approach

The clearest limitation of this study is that we tested STORM using computational models and have yet to deploy it on real neurons. We have started with a modeling framework to demonstrate that STORM is conceptually and computationally feasible, which was not obvious a priori given that it was not clear if natural movies would contain identifiable stimulus directions that differentially affect responses of target and non-target cell populations. However, the success of the approach does depend significantly on model accuracy (Fig. 6), which current best models fall just shy of achieving, particularly when generalizing across retinas and naturalistic stimulus conditions [Shah et al., 2022, Idrees et al., 2024, D’Agostino et al., 2025]. At the same time, we and others are making rapid progress toward models that are sufficiently accurate to serve as a proxy for the real neuron and models that generalize across retinas not included in training, with such models becoming available [Idrees et al., 2026]. Furthermore, hybrid bio-CNN models that include more biological detail and thus generalize better across conditions are also being actively developed [Idrees et al., 2024]. Thus, we plan to test these approaches directly as increasingly accurate models become available.

As hinted at above, stimuli optimized to be metameric using gradients from a model fit to data from a single retina may not remain metameric when applied to a different retina, since model fits—and the underlying gradients they produce—can vary across individuals. One strategy to improve robustness to this variability would be to use an ensemble of models—one fit to each of several retinas— to compute gradients for the non-target cells rather than relying on gradients from a single fitted model. Stimulus directions can be restricted to those that remain orthogonal to the non-target gradients across all models in the ensemble, not just one. This approach would favor directions that are robust to model-fitting variability, rather than directions that would be metameric only for a single model’s idiosyncrasies. We leave an in-depth investigation of these questions to an upcoming study.

The translation-invariant approach, though key to making this method practical, assumes a degree of homogeneity within each cell type. Given that the approach was robust to some model inaccuracies, there is hope that modest cell-to-cell heterogeneity will not prevent the approach from working. However, it will likely fail if within-type variability becomes comparable to the differences between types. In primate retina, functional clustering studies indicate that this is not the case: RGCs form well-separated, near-homogeneous clusters corresponding to irreducible cell types [Field et al., 2007], though this may not hold in other sensory modalities such as olfaction [Araneda et al., 2000]. It may also fail in cases where there are substantial changes in the response properties between cell types across retinal eccentricity (correspondingly across the visual field). Indeed, the receptive field size and temporal integration of midget and parasol RGCs changes substantial between foveal and peripheral vision. Thus, applications of this approach to behaving animals will likely need to focus on a particular location in the visual field.

Another significant limitation is the focus on just four RGC types. While these are the numerically dominant RGC types of the primate retina [Field and Chichilnisky, 2007], the primate retina contains at least 17 cell types [Dacey et al., 2003, Peng et al., 2019], and likely many more [Yamada et al., 2005, Masri et al., 2019, Kling et al., 2024]. The mouse retina contains ~45 RGC types [Goetz et al., 2022]. As more cell types are added, so are more constraints, which could challenge the STORM approach. However, many of these RGC types exhibit quite nonlinear and/or highly selective responses. These nonlinear responses may significantly mitigate the impact of these additional types on generating cell-type specific stimuli (see previous subsection). Nevertheless, generating stimuli that are genuinely metameric to 44 of the 45 RGC types in the rodent retina will very likely require accurate models of all 45 types.

The approach also has several technical considerations. For example, direct orthogonalization against the sensitivity of every non-target RGC at every spatial location and time point becomes computationally expensive. To address this challenge, we used randomized SVD to identify a low-rank approximation of the non-target sensitivity subspace (see Methods; Supplementary Fig. S15). The low-rank SVD approach makes selective manipulation in this high-dimensional stimulus space much more computationally feasible. Additionally, there is the question of how to choose the ‘original stimulus’ that will be modified by STORM. This choice can have some critical outcomes. For example, if a certain cell type is completely unresponsive to the stimulus, it will be challenging to manipulate the activity of this cell type. Conversely, selecting a base movie to which non-target cells are relatively insensitive may facilitate the production of a stimulus that is metameric for the non-target cells. Thus, some starting stimuli may facilitate or inhibit the success of the approach. We plan to explore this issue more in future work.

### Future applications of cell-type-specific manipulations

An important future application of this approach is in ophthalmic diagnostics. Notably, the 20-30 RGC types within the primate retina appear to have differential levels of vulnerability to degenerative diseases like glaucoma [Karamanlis et al., 2022, Quigley, 1999], retinitis pigmentosa [Dyszkant et al., 2025] and diabetic retinopathy [Cui et al., 2019]. This means that vision tests that isolate the function of individual RGC types may be able to diagnose these diseases at their earliest stages, when only the most-vulnerable RGC types have been affected, but before more profound vision loss (involving more RGC types) has occurred. Such an approach may allow treatment to begin earlier, thereby slowing down degeneration and retaining visual function for longer.

This differential vulnerability in retinal function may extend to neurodegenerative diseases such as Alzheimer’s, Parkinson’s, Leber’s and Huntington’s disease [La Morgia et al., 2017b,a, Kersten et al., 2015], suggesting RGC type-specific vision tests could offer an early biomarker for these conditions as well. Beyond diagnostics, RGC type-specific stimulation may also have therapeutic potential in conditions associated with progressive vision loss by preserving RGC function and slowing degeneration. RGC survival depends on BDNF-TrkB neurotrophic signaling which is supported by neural activity. In optic neuropathies, impaired retrograde transport of BDNF and reduced RGC activity can weaken this trophic support and contribute to progressive degeneration. Restoring physiological activity selectively in vulnerable RGC types may therefore help maintain responsiveness to neurotrophic support and preserve cell survival pathways [Lambuk et al., 2022].

Moreover, understanding how each RGC type encodes visual information is crucial for development of vision restoration technologies such as retinal prosthetics. Current implants do not account for functional diversity of RGC types and the specific visual features each type encodes [Mills et al., 2017, Ayton et al., 2020]. Understanding what each RGC type encodes could significantly improve retinal prostheses by enabling selective stimulation of specific RGC types rather than indiscriminately activating all cells near an electrode. Recent work has shown that selective electrical stimulation of RGCs can faithfully reproduce naturalistic retinal activity [Phillips et al., 2026]. Such approaches could enhance the quality and fidelity of restored vision.

## Methods

### Linear-Nonlinear Poisson (LNP) RGC models

For the results in Fig. 2, we used the Linear-Nonlinear Poisson (LNP) model which is a well-established framework for modeling retinal ganglion cell (RGC) responses. Using literature-derived parameters for receptive field (RF) size, temporal dynamics, and nonlinear response properties [Kling et al., 2020, Parthasarathy et al., 2017], we simulated the responses of four major RGC types in primate: ON/OFF parasol and ON/OFF midget.

Spatial receptive fields were modeled as Difference-of-Gaussian filters in which centers represent the excitatory region while the surrounds capture the antagonistic region of the receptive field. Each RGC type forms its own mosaic tiling the visual field. This mosaic arrangement ensures that every region of the visual field is represented consistently, minimizing the gaps and redundancy in visual signaling to the brain [Wassle and Boycott, 1991, Wässle et al., 1981, Devries and Baylor, 1997].

For Gaussian function

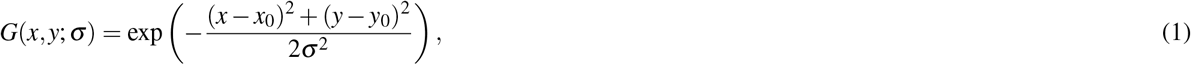

with center and surround components normalized as

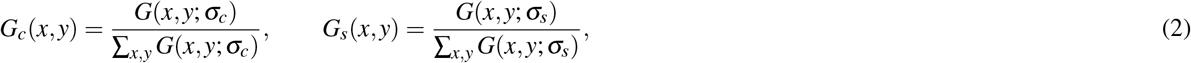

the spatial receptive fields were

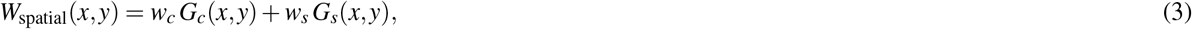

where *σ*_*c*_, *σ*_*s*_ and *w*_*c*_, *w*_*s*_ set RF size and center-surround balance.

Temporal filters were defined using biphasic filters with a primary lobe indicating polarity (ON vs OFF) and a secondary lobe setting response transience and integration time. We first defined two temporal Gaussians:

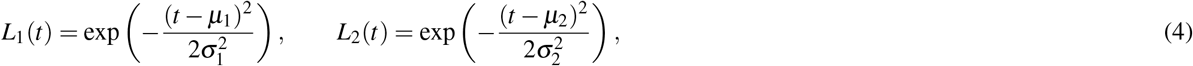

which were then combined to form the biphasic temporal filters *h*(*t*) for ON and OFF cells:

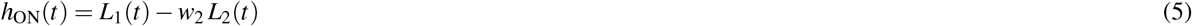

and

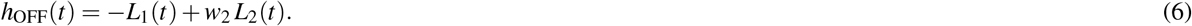

The parameter *w*_2_ defines the relative magnitude of the negative or positive lobe of the receptive field, for ON and OFF cells, respectively.

Combining these temporal filters with the spatial receptive fields, the separable spatiotemporal filter was obtained for each neuron:

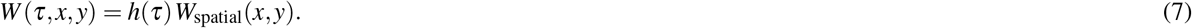

The parameters *σ*_*c*_, *σ*_*s*_ and *w*_*c*_, *w*_*s*_, *w*_2_, *σ*_1_, and *σ*_2_ were chosen separately for each cell type such that the model RGC receptive fields resembled experimental observations[Kling et al., 2020, Parthasarathy et al., 2017].

These spatiotemporal receptive fields were then normalized to have unit Euclidean norm:

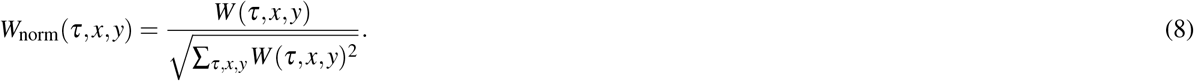

In the LNP model, the stimulus is first passed through the same spatiotemporal filter which produces a generator signal that represents a weighted sum of the stimulus intensities over space and time:

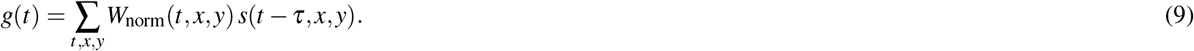

The generator signal is then passed through an instantaneous nonlinear function to introduce nonlinearities such as spike threshold and response saturation to transform the signal into the spike rate. This step models the neural response to the stimulus:

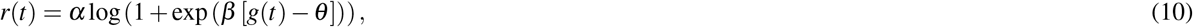

where *α* is a gain factor, *β* sets the slope of the nonlinearity, and *θ* is a threshold parameter. Threshold, gain, and saturation values were chosen so that the models would approximate experimental measurements [Parthasarathy et al., 2017].

Finally, spike counts were determined by a Poisson process that models the random nature of spike generation with the spike rate serving as the rate parameter for the Poisson process [Chichilnisky, 2001]:

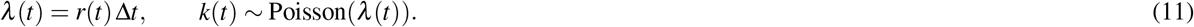

### LN-LN cascade RGC models

For the results in Fig. 3, RGC responses were modeled using an LN-LN cascade to capture subunit nonlinearities [Maheswaranathan et al., 2018, Wu et al., 2015]. At each pixel location (*x, y*), the stimulus is convolved with a temporal filter *h*(*τ*) to produce a generator signal g(t,x,y),

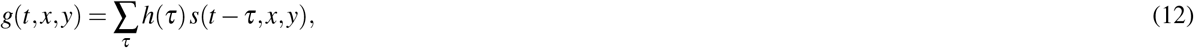

where *h*(*τ*) is the biphasic temporal filter from Eq. 5 or 6, depending on the RGC polarity (ON vs OFF). Subunit nonlinearity was then applied independently at each location:

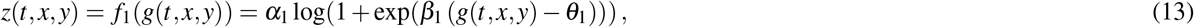

where *α*_1_, *β*_1_, and *θ*_1_ are the subunit gain, slope, and threshold parameters.

The nonlinearly transformed pixel outputs are linearly combined, weighted the spatial receptive field:

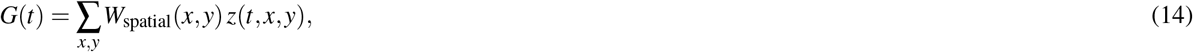

where *W*_spatial_(*x, y*) is the difference-of-Gaussians spatial filter defined by Eq. 3.

This pooled signal was passed through a static nonlinearity to produce the instantaneous firing rate:

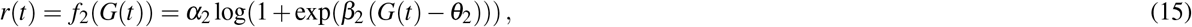

where *α*_2_, *β*_2_, and *θ*_2_ are the output-stage gain, slope, and threshold parameters.

Similar to the LN model, the model parameters were chosen separately for each RGC type, consistent with prior studies [Maheswaranathan et al., 2018, Wu et al., 2015].

### Primate retina data for training CNN models

We used the electrophysiological dataset from a previous study [Idrees et al., 2024]. Recordings were obtained from an ex vivo macaque retina using a high density multielectrode array. RGC activity was recorded during 36 minutes of binary checkerboard white noise and 9 minutes of grayscale naturalistic movie presentation at a mean light level of 50 R*receptor^−1^s^−1^. The checkerboard stimulus consisted of a 100×75 pixel grid with each pixel corresponding to 30 *µ*m on the retinal surface. The naturalistic movies were generated by displacing natural scene images from the Van Hateren dataset [Van Hateren and van der Schaaf, 1998] using eye-movement trajectories derived from the DOVES dataset [Van Der Linde et al., 2009]. We selected 84 RGCs (16 ON and 13 OFF parasol cells and 40 ON and 15 OFF midget cells) for modeling purposes based on spike sorting quality and reliability across experimental conditions.

### CNN model with fully-connected output layer

For the results in Fig. 4, we used a CNN model described in our previous study [Idrees et al., 2024] based on an architecture similar to Deep Retina [McIntosh et al., 2016]. The model comprises three convolutional layers followed by a fully-connected layer, with 10, 15 and 20 channels and 11×11, 7×7 and 7×7 spatial filters respectively. (For analyses related to translation invariance, we used a different model with no fully-connected layer: that model is described in Section CNN model with firing rate map output). The CNN model processes spatiotemporal movie segments (80 frames, 8.3 ms per frame) using spatiotemporal convolutional layers, followed by normalization, max-pooling, and a fully connected readout with trainable output activation functions. The network was trained to predict the instantaneous firing rates of primate RGCs from binary checkerboard white-noise stimuli using the Adam optimizer[Kingma and Ba, 2014] and a Poisson negative log-likelihood loss. Regularization consisted of *L*_2_ weight penalties together with an *L*_1_ penalty on the readout layer to obtain sparse feature representations. Learning rate was set to 0.001.

### Synthetic CNN-based RGC mosaics

After spike sorting, gaps may appear within an RGC mosaic and they do not necessarily indicate missing RGCs. They represent areas where cells were missed during spike sorting or where the recording technology failed.

The CNN model described in the preceding section (and in Fig. 4A) was trained only on 84 RGCs that passed our data-quality criteria so the mosaic contained such gaps. This is a limitation when evaluating selective silencing because fewer RGCs constrain the stimulus which increases the degrees of freedom and makes selective silencing artificially easier. To demonstrate that the method of tailored stimulus modification can selectively target every cell under more realistic and fully constrained conditions, we constructed a complete synthetic mosaic (Fig. 4B) using the CNN model described in the preceding section. The model was trained on white noise checkerboard and naturalistic movie stimuli. The CNN learns a set of weights for each model RGC, and additional “synthetic” RGCs were generated by spatially shifting the stimulus relative to an existing model RGC receptive field to predict the responses of RGCs at different locations. Shifting the stimulus by(*x*_0_ −*v*_*x*_,; *y*_0_ −*v*_*y*_) relocates the response of a cell with receptive field center (*x*_0_, *y*_0_) to the target location (*v*_*x*_, *v*_*y*_).

After translation, the stimulus was padded with gray pixels of mean stimulus intensity to reduce edge artifacts. Synthetic cells were arranged on a regular grid covering the visual field to create a complete mosaic. A 20-pixel margin was excluded around the edges of the visual field so every retained cell was far enough from the border for its receptive field to stay entirely over the real stimulus and to never overlap the blank regions created by padding the shifted stimulus. Grid spacing was set separately for each RGC type to match the center-to-center spacing of the recorded RGCs: 11 pixels for parasol cells and 7 pixels for midget cells, corresponding to approximately 330 um and 210 um on the retina (30 um per checker). We evaluated two mosaic configurations: homogeneous and heterogeneous.

In the homogeneous mosaic (Fig. 4C), all cells of a given type were generated from spatial translations of a single template cell. For each type, the template was chosen from the cells within the border with high predictive performance. Synthetic cells were generated by translating the stimulus so that the receptive field of template was centered at each grid location. As a result, all cells of a given type shared identical spatial, temporal, and nonlinear response properties, differing only in spatial receptive field location.

In the heterogeneous mosaic (Fig. S8 C), a complete mosaic was generated from multiple experimentally recorded cells of each RGC type to better capture biological variability within each cell type. Cells with receptive field centers located within the ROI (excluding the 20-pixel border) were retained at their original locations and remaining gaps were filled by randomly selecting recorded RGCs and translating their receptive fields to the vacant positions. A synthetic cell was placed only if its target location was farther than the specified spacing from every previously placed cell to maintain physiologically realistic spacing between neighboring receptive fields. The resulting mosaic preserved natural variability in receptive field size, temporal dynamics, and output nonlinearity within each type.

### CNN model with firing rate map output

For the results in Fig. 5 we used a spatially translation-invariant version of the CNN model from Section CNN model with fully-connected output layer, the final fully connected layer was replaced with a 1×1 convolution with four output channels, one per RGC type. The model has three convolutional layers with 10, 15, and 20 channels and 11 ×11, 7× 7, and 7× 7 spatial filters, and all layers use padding to preserve spatial dimensions throughout the network. The 1×1 convolution applies a learned linear combination of the 20 feature channels independently at each spatial location, producing four firing rate maps, one per RGC type, over the full spatial extent of the stimulus. This is equivalent to applying the same feature readout weights at every spatial position, enforcing translation invariance in the mapping from features to firing rates.

The model was trained to minimize the mean absolute error between predicted and observed firing rates. Learning rate was set to 0.0001.The predicted firing rate for each cell was computed by averaging the firing rate map over a 3×3 pixel grid centered at the cell’s receptive field center, and this spatially averaged prediction was matched against the cell’s observed firing rate. This formulation allowed the model to be tolerant to small errors in receptive field center localization.

### SpatioTemporal Orthogonal Response Modulation (STORM)

This section describes the SpatioTemporal Orthogonal Response Modulation (STORM) procedure for constructing a stimulus movie that selectively modulates the responses of a target population of RGCs while leaving the responses of non-target RGCs unchanged. The key quantity is the gradient of each neuron’s predicted firing rate with respect to the stimulus movie — a vector in stimulus space that points in the direction along which that neuron’s response increases most steeply. By computing these gradients for both target and non-target neurons, the stimulus space can be decomposed into two subspaces: one spanned by the directions that drive target neurons, and one spanned by the directions that drive non-target neurons. A stimulus modification that lies entirely within the target subspace but is orthogonal to the non-target subspace will modulate target responses while leaving non-target responses unchanged by construction (Fig1). Applying such modifications to a movie **s** yields a modified movie **s**\* — the two together forming a metameric pair with respect to the non-target population: they produce equivalent responses in non-target RGCs while differing systematically in their drive to target RGCs. We begin with initializing the modified stimulus movie from the original movie, **s**\* = **s** ∈ ℝ^*F*×*H*×*W*^ of *F* frames, *H* height, and *W* width, and update it iteratively:

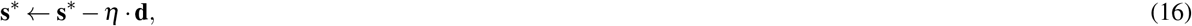

where *η* is the step size, chosen large enough to produce a substantial change in target neuron firing rates while small enough to induce minimal changes in non-target responses, and **d** ∈ℝ^*F*×*H*×*W*^ is the stimulus update direction that points toward increasing target neuron responses while remaining orthogonal to the non-target subspace. Subtracting *η* **d** moves the stimulus in the direction of steepest decrease in target responses while leaving non-target responses unchanged by construction. The remainder of this section describes how **d** is computed.

The models do not process the full stimulus movie **s**\*, in a single pass. Instead, they operate on temporal windows of fixed length *T*, where each window *w* corresponds to a *T*-frame except **s**_*w*:*w*+*T*_ ∈ ℝ^*T* ×*H*×*W*^ representing the recent stimulus history available to the retina at time *w*. These windows are formed by rolling **s**\* with a stride of 1, yielding *N* = *F*− *T* + 1 overlapping windows that together span the entire movie. Since adjacent windows share *T*− 1 frames, the same frame *f* appears in up to *T* different windows simultaneously — a modification to frame *f* therefore affects the responses of all windows that contain it. The natural space for stimulus optimization is thus ℝ^*F*×*H*×*W*^ rather than ℝ^*N*×*T* ×*H*×*W*^, since the latter would assign a separate gradient to frame *f* for each window it appears in, leaving no consistent answer to how frame *f* should actually be modified. For each modeled neuron *i* and temporal window *w*, the gradient of its predicted firing rate *r*_*i,w*_ with respect to the *T*-frame input has shape ℝ^*T* ×*H*×*W*^ and is embedded into the full movie space by placing it at temporal position [*w* : *w* + *T*], yielding a vector

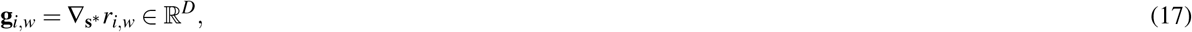

where *D* = *F*× *H* ×*W*, that captures how a modification to any frame of the full movie affects neuron *i*’s response at window *w*. Because this gradient depends on which *T* frames are currently in the input window, it is computed separately for each of the *N* temporal windows, yielding a distinct gradient vector **g**_*i,w*_ per (neuron, window) pair. Stacking these vectors as rows over all neurons and windows yields the gradient matrix

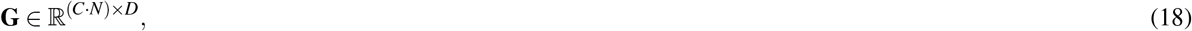

where *C* is the total number of modeled neurons. The rows of **G** span the subspace defining the set of stimulus directions that collectively drive the modeled population of *C* neurons across all *N* temporal windows. Neurons are then partitioned into two sets: *target neurons* whose responses are to be suppressed or enhanced, giving rise to 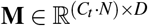, and *non-target neurons* whose responses are to be held fixed, giving rise to 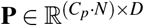, where *C*_*t*_ and *C*_*p*_ are the number of target and non-target neurons respectively.

The rows of **P** collectively span the *non-target subspace* in ℝ^*D*^ — the set of stimulus directions that drive non-target neurons. Although **P** may contain a large number of rows, the effective dimensionality of this subspace is typically much lower than *C*_*p*_ ·*N*, since gradient vectors from nearby neurons and overlapping temporal windows tend to be highly correlated. An effective stimulus modification must remain orthogonal to this subspace, meaning that the stimulus update direction must be perpendicular to every vector in the non-target subspace. This is achieved in two steps: first an orthonormal basis for the non-target subspace is constructed from the rows of **P**, and then the target gradients in **M** are orthogonalized against this basis by removing any component that lies within the non-target subspace.

For the first step of constructing an orthonormal basis, a set of vectors {**q**_1_, **q**_2_, …, **q**_*k*_} in ℝ^*D*^ is sought that spans the same subspace as the rows of **P**, where each basis vector has unit length (∥**q** _*j*_∥ = 1) and any two distinct basis vectors are perpendicular (**q** _*j*_ · **q**_*l*_ = 0 for *j* ≠ *l*). This basis is obtained by applying a singular value decomposition to **P**:

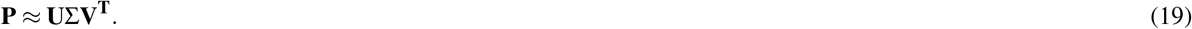

The right singular vectors — the rows of *V*^*T*^ — are guaranteed to be orthonormal by construction, and represent the principal directions of the non-target subspace ordered by the amount of variance they capture across the rows of **P**. Each such vector represents a stimulus pattern that produces a coordinated response across the population of non-target neurons. Because **P** can be very large — with *C*_*p*_ ·*N* rows each of dimension *D* — the randomized SVD algorithm [Halko et al., 2011] is applied to rows of **P**, which computes an accurate low-rank approximation at a fraction of the cost of full SVD by first compressing the matrix via random projection before factorization. Retaining the top *k* right singular vectors yields *Q* = *V*^*T*^ ∈ ℝ^*k*×*D*^. The rank *k* was determined empirically by running randomized SVD once with a conservatively large number of components and inspecting the cumulative variance explained by the singular value spectrum; *k* was then set to the smallest number of components that account for at least 99% of the total variance in **P**. For analysis presented in this manuscript, 53 *< k <* 687 with k varying depending on the movie length, number of RGCs, and number of model parameters.

For the second step, the projection of **M** onto the non-target subspace spanned by **Q** is removed:

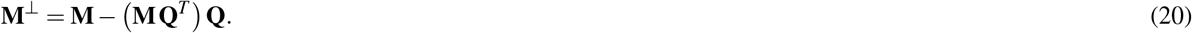

The residual matrix **M**^⊥^ contains rows that drive target neurons while remaining orthogonal to the non-target subspace by construction, i.e. **M**^⊥^**Q**^*T*^ = **0**. Denoting each row of **M**^⊥^ corresponding to neuron *i* and window *w* as 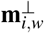, the stimulus update direction **d** is obtained by summing over all *N* temporal windows and averaging over *C*_*t*_ target neurons:

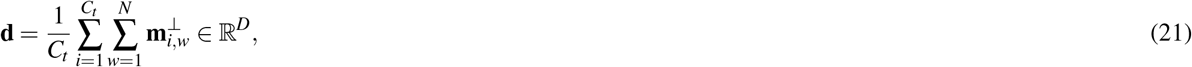

reshaped back to ℝ^*F*×*H*×*W*^.

The stimulus update direction **d** is normalized to unit length,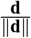, before applying the update **s**\* ← **s**\* − *η* · **d**, so that the step size *η*, controls the magnitude of the stimulus modification independently of the number of target neurons or temporal windows.

At each iteration, **s**\*← **s**\* −*η* ·**d**, the gradient matrices **M** and **P** are recomputed on the current **s**\* and a new orthonormal basis **Q** is constructed. Recomputing gradients at each iteration is necessary because the optimal modification direction depends on the current stimulus — as the stimulus changes, it can engage different nonlinearities in the circuit, and the gradient directions can change. Thus the procedure is applied iteratively such that at each step only the target cells are modulated while leaving the non-target cells unaffected. Iterations continue until the mean predicted firing rate of all target neurons, averaged across all *N* temporal windows, falls below a threshold of 0.001. The stimulus at convergence is taken as **s**\*.

The stimuli used to train and test the models spanned the full intensity range of the display, such that perturbations of pixels near the range limits could produce modified stimulus values outside the displayable range. For experimental implementation, a narrower baseline stimulus range would be used to allow sufficient room for modification, with additional constraints to keep the final stimuli within the displayable range.

In the case of models with a fully connected readout layer (Fig 4), individual modeled neurons correspond directly to identified RGCs, and target and non-target sets are defined by selecting specific cells. In the case of the translation-invariant model (Fig 5), the output is a firing-rate map — one per RGC type — and target and non-target sets are instead defined by sampling spatial locations within these maps, with each sampled location contributing one gradient vector per temporal window. For CNN-based models, gradients are computed via reverse-mode automatic differentiation using JAX Python library, which efficiently propagates the gradient from a scalar output back to every stimulus pixel in a single backward pass. For simpler models — linear, LN, and LN-LN (Figs 2,3) — gradients are computed analytically, since the functional form of these models admits closed-form expressions for ∇_**s**\*_*r*_*i,w*_.

### Gradient computation for the Linear, LNP and LN-LN cascade Models

For the LNP and LN-LN cascade used in Figs 2 and 3, gradients were computed analytically. In the linear model, each neuron’s response is given by the generator signal:

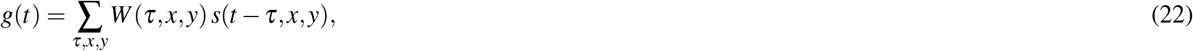

where *s*(*t, x, y*) is the stimulus intensity at time *t* and pixel (*x, y*), and *W* (*τ, x, y*) is the neuron’s spatiotemporal RF. The model RGC response is:

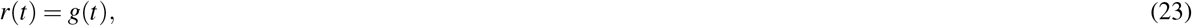

such the gradient of response with respect to the stimulus equals the RF:

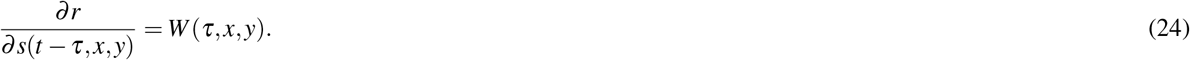

In the Linear-Nonlinear-Poisson (LNP) model, the same linear stage is followed by a static nonlinearity and a Poisson spike generator. We used a softplus nonlinearity:

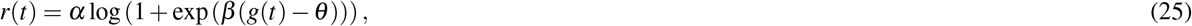

where *α* is a gain, *β* a sensitivity, and *θ* a threshold parameter. The gradient factorizes via the chain rule as:

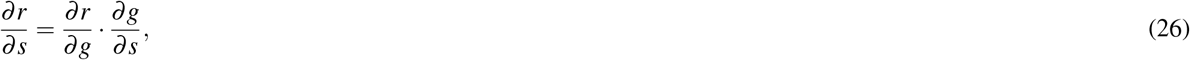

with

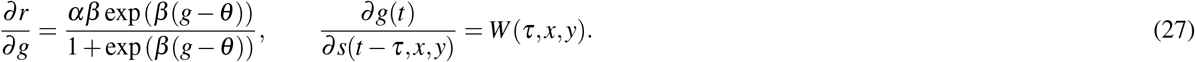

The Poisson spike count for a given bin is then drawn with rate parameter *r*(*t*), but spike stochasticity doesn’t affect the gradient used for stimulus optimization.

In the LN-LN cascade model, the gradient of the firing rate with respect to the stimulus is again obtained by applying the chain rule:

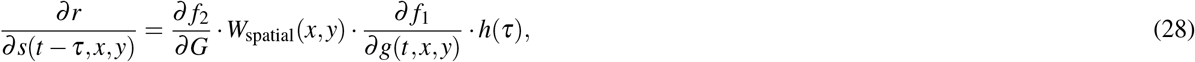

with

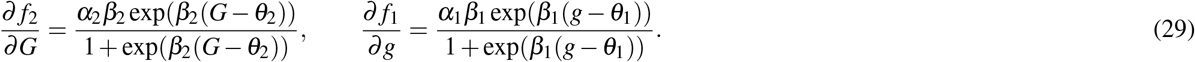

### Robustness Test

STORM relies on an accurate encoding model to estimate the gradients used to suppress one RGC type while remaining metameric to the others. To assess the sensitivity of the method to model error (Fig. 6), we trained the map-based CNN model described in Section CNN model with firing rate map output for 20 epochs and saved multiple checkpoints throughout the training. Checkpoints from earlier stages of training were treated as imperfect models and used to generate the modified stimuli whereas the checkpoint with the highest predictive performance was used as the tester model (Fig. 6A). All models shared the same architecture and differed only in the amount of training.

Similarity between a generator and the tester was evaluated as the Pearson correlation between their predicted responses to the same held-out stimuli, computed as:

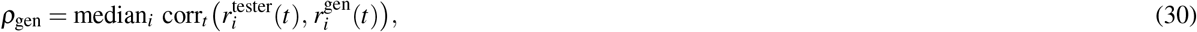

Here 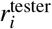and 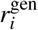are the responses predicted by the tester and generator models to the same stimuli and corr_*t*_ denotes the correlation over time.

For the target RGC, the relative response reduction predicted by the tester and generator models is,

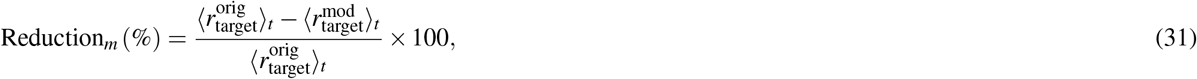

where *r*^orig^ and *r*^mod^ are the responses predicted by model *m* to the original and modified movies. From these, we report the fraction of the suppression designed by the generator that transferred to the tester(Fig. 6E):

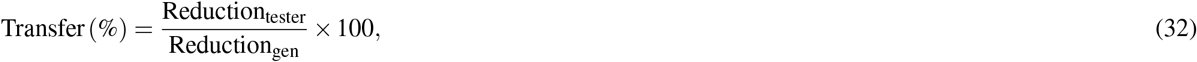

For the non-target RGC, responses may change in either direction, so we report the absolute deviation over time relative to the original firing rate (Fig. 6F):

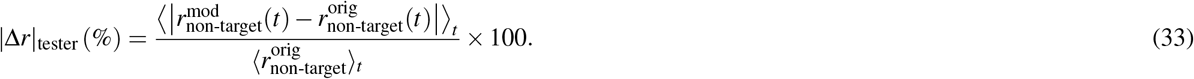

where *r*^orig^ and *r*^mod^ denote the responses predicted by the tester model to the original and modified stimuli respectively.

## Acknowledgements

This work was supported by Grant # R01-EY036594 (GDF, FR, and JZ) and # P30-EY000331 (GDF, JZ) from the National Institutes of Health, an Unrestricted grant from Research to Prevent Blindness to the Jules Stein Eye Institute (GDF, JZ), Grant # FA9550-26-1-B113 from the Air Force Office of Scientific Research (JZ), a CIFAR Fellow Award for Learning in Machines and Brains (JZ), the Joan and Jerome Snyder Chair in Vision Science at UCLA (JZ), and the Fran and Ray Stark Chair in Ophthalmology (GDF). We thank Huakun Li for valuable discussions, thoughtful feedback, and helpful suggestions.

## Author contributions statement

N.G., S.I., V.R., F.R., G.F., and J.Z. conceptualized the experiments. N.G. and S.I. conducted the experiments. N.G. and S.I. created the visualizations. M.M., F.R., G.F., and J.Z. supervised the work. F.R., G.F., and J.Z. secured the funding. All authors reviewed the manuscript.

## References

Ricardo C Araneda, Abhay D Kini, and Stuart Firestein. The molecular receptive range of an odorant receptor. Nature neuroscience, 3(12):1248–1255, 2000.

Blaine N Armbruster, Xiang Li, Mark H Pausch, Stefan Herlitze, and Bryan L Roth. Evolving the lock to fit the key to create a family of g protein-coupled receptors potently activated by an inert ligand. Proceedings of the National Academy of Sciences, 104(12):5163–5168, 2007.

Lauren N Ayton, Nick Barnes, Gislin Dagnelie, Takashi Fujikado, Georges Goetz, Ralf Hornig, Bryan W Jones, Mahiul MK Muqit, Daniel L Rathbun, Katarina Stingl, et al. An update on retinal prostheses. Clinical Neurophysiology, 131(6):1383–1398, 2020.

Stephen A Baccus and Markus Meister. Fast and slow contrast adaptation in retinal circuitry. Neuron, 36(5):909–919, 2002.

Pouya Bashivan, Kohitij Kar, and James J DiCarlo. Neural population control via deep image synthesis. Science, 364(6439):eaav9436, 2019.

Michael J Berry, Iman H Brivanlou, Thomas A Jordan, and Markus Meister. Anticipation of moving stimuli by the retina. Nature, 398(6725):334–338, 1999.

Edward S Boyden, Feng Zhang, Ernst Bamberg, Georg Nagel, and Karl Deisseroth. Millisecond-timescale, genetically targeted optical control of neural activity. Nature neuroscience, 8(9):1263–1268, 2005.

Max F Burg, Thomas Zenkel, Michaela Vystrčilová, Jonathan Oesterle, Larissa Höfling, Konstantin F Willeke, Jan Lause, Sarah Müller, Paul Fahey, Zhiwei Ding, et al. Most discriminative stimuli for functional cell type clustering. In International Conference on Learning Representations, volume 2024, pages 22508–22533, 2024.

Jon Cafaro and Fred Rieke. Regulation of spatial selectivity by crossover inhibition. Journal of Neuroscience, 33(15):6310–6320, 2013.

EJ Chichilnisky. A simple white noise analysis of neuronal lightresponses. Network: computation in neural systems, 12(2):199, 2001.

Run-Ze Cui, Lu Wang, Sheng-Nan Qiao, Yong-Chen Wang, Xin Wang, Fei Yuan, Shi-Jun Weng, Xiong-Li Yang, and Yong-Mei Zhong. On-type retinal ganglion cells are preferentially affected in stz-induced diabetic mice. Investigative Ophthalmology & Visual Science, 60(5):1644–1656, 2019.

Dennis M Dacey. Origins of perception: retinal ganglion cell diversity and the creation of parallel visual pathways. In Michael S Gazzaniga, editor, The Cognitive Neurosciences, pages 281–301. MIT Press, Cambridge, MA, 2004.

Dennis M Dacey, Beth B Peterson, Farrel R Robinson, and Paul D Gamlin. Fireworks in the primate retina: in vitro photodynamics reveals diverse lgn-projecting ganglion cell types. Neuron, 37(1):15–27, 2003.

Andrew M Derrington, John Krauskopf, and Peter Lennie. Chromatic mechanisms in lateral geniculate nucleus of macaque. The Journal of physiology, 357(1):241–265, 1984.

Steven H Devries and Denis A Baylor. Mosaic arrangement of ganglion cell receptive fields in rabbit retina. Journal of neurophysiology, 78(4):2048–2060, 1997.

Nadine Dyszkant, Jonathan Oesterle, Yongrong Qiu, Merle Harrer, Timm Schubert, Dominic Gonschorek, and Thomas Euler. Photoreceptor degeneration has heterogeneous effects on functional retinal ganglion cell types. The Journal of Physiology, 603(21):6599–6621, 2025.

Federico D’Agostino, Thomas Zenkel, Baptiste Lorenzi, Michaela Vystrčilová, Dominic Gonschorek, Samuel Suhai, Samuele Virgili, Alexander S Ecker, Olivier Marre, Larissa Höfling, et al. The openretina project: Collaborative retina modelling across datasets and species. bioRxiv, pages 2025–03, 2025.

Christina Enroth-Cugell and John G Robson. The contrast sensitivity of retinal ganglion cells of the cat. The Journal of physiology, 187(3):517–552, 1966.

O Estévez and Henk Spekreijse. The “silent substitution” method in visual research. Vision research, 22(6):681–691, 1982.

Kayson Fakhar, Shrey Dixit, Fatemeh Hadaeghi, Konrad P Kording, and Claus C Hilgetag. When neural activity fails to reveal causal contributions. bioRxiv, 2023.

Lief Fenno, Ofer Yizhar, and Karl Deisseroth. The development and application of optogenetics. Annual review of neuroscience, 34(2011):389–412, 2011.

Greg D Field and EJ Chichilnisky. Information processing in the primate retina: circuitry and coding. Annu. Rev. Neurosci., 30 (1):1–30, 2007.

Greg D Field, Alexander Sher, Jeffrey L Gauthier, Martin Greschner, Jonathon Shlens, Alan M Litke, and EJ Chichilnisky. Spatial properties and functional organization of small bistratified ganglion cells in primate retina. Journal of Neuroscience, 27(48):13261–13272, 2007.

Jeremy Freeman, Greg D Field, Peter H Li, Martin Greschner, Deborah E Gunning, Keith Mathieson, Alexander Sher, Alan M Litke, Liam Paninski, Eero P Simoncelli, et al. Mapping nonlinear receptive field structure in primate retina at single cone resolution. Elife, 4:e05241, 2015.

Jeffrey L Gauthier, Greg D Field, Alexander Sher, Martin Greschner, Jonathon Shlens, Alan M Litke, and EJ Chichilnisky. Receptive fields in primate retina are coordinated to sample visual space more uniformly. PLoS biology, 7(4):e1000063, 2009.

Guy Gaziv, Sarah Goulding, Ani Ayvazian-Hancock, Yoon Bai, and James J DiCarlo. Noninvasive precision modulation of high-level neural population activity via natural vision perturbations. arXiv preprint 2506.05633, 2025.

Jillian Goetz, Zachary F Jessen, Anne Jacobi, Adam Mani, Sam Cooler, Devon Greer, Sabah Kadri, Jeremy Segal, Karthik Shekhar, Joshua R Sanes, et al. Unified classification of mouse retinal ganglion cells using function, morphology, and gene expression. Cell reports, 40(2), 2022.

Alex R Gogliettino, Sam Cooler, Ramandeep S Vilkhu, Nora J Brackbill, Colleen Rhoades, Eric G Wu, Alexandra Kling, Alexander Sher, Alan M Litke, and EJ Chichilnisky. Modeling responses of macaque and human retinal ganglion cells to natural images using a convolutional neural network. bioRxiv, pages 2024–03, 2024.

Tim Gollisch and Markus Meister. Eye smarter than scientists believed: neural computations in circuits of the retina. Neuron, 65(2):150–164, 2010.

Nathan Halko, Per-Gunnar Martinsson, and Joel A Tropp. Finding structure with randomness: Probabilistic algorithms for constructing approximate matrix decompositions. SIAM review, 53(2):217–288, 2011.

S Hochstein and RM Shapley. Quantitative analysis of retinal ganglion cell classifications. The Journal of physiology, 262(2):237–264, 1976.

Larissa Höfling, Klaudia P Szatko, Christian Behrens, Yuyao Deng, Yongrong Qiu, David Alexander Klindt, Zachary Jessen, Gregory W Schwartz, Matthias Bethge, Philipp Berens, Katrin Franke, Alexander S Ecker, and Thomas Euler. A chromatic feature detector in the retina signals visual context changes. eLife, 13, 10 2024. doi: 10.7554/ELIFE.86860.

Saad Idrees, Michael B Manookin, Fred Rieke, Greg D Field, and Joel Zylberberg. Biophysical neural adaptation mechanisms enable artificial neural networks to capture dynamic retinal computation. Nature Communications, 15(1):5957, 2024.

Saad Idrees, Bella Hofflich, Alexandra Kling, Alexander Sher, Alan Litke, E.J. Chichilnisky, and Joel Zylberberg. Minimalist CNN models explain primate midget and parasol RGC responses with accuracy near the achievable limit. Poster presented at the FASEB Retinal Neurobiology and Visual Processing Conference, Sacramento, CA, U.S.A., 2026.

Makoto Ishihara. Versuch einer deutung der photoelektrischen schwankungen am froschauge. Archiv für die gesamte Physiologie des Menschen und der Tiere, 114(11):569–618, 1906.

Dimokratis Karamanlis, Helene Marianne Schreyer, and Tim Gollisch. Retinal encoding of natural scenes. Annual Review of Vision Science, 8:171–193, 2022.

Hannah M Kersten, Helen V Danesh-Meyer, Dean H Kilfoyle, and Richard H Roxburgh. Optical coherence tomography findings in huntington’s disease: a potential biomarker of disease progression. Journal of Neurology, 262(11):2457–2465, 2015.

Kerry J Kim and Fred Rieke. Temporal contrast adaptation in the input and output signals of salamander retinal ganglion cells. The Journal of Neuroscience, 21(1):287–299, 2001.

Diederik P Kingma and Jimmy Ba. Adam: A method for stochastic optimization. arXiv preprint 1412.6980, 2014.

A Kling, S Cooler, MB Manookin, C Rhoades, N Brackbill, G Field, F Rieke, A Sher, A Litke, and EJ Chichilnisky. Functional diversity in the output of the primate retina. BioRxiv, pages 2024–10, 2024.

Alexandra Kling, Alex R Gogliettino, Nishal P Shah, Eric G Wu, Nora Brackbill, Alexander Sher, Alan M Litke, Ruwan A Silva, and EJ Chichilnisky. Functional organization of midget and parasol ganglion cells in the human retina. BioRxiv, pages 2020–08, 2020.

Chiara La Morgia, Lidia Di Vito, Valerio Carelli, and Michele Carbonelli. Patterns of retinal ganglion cell damage in neurodegenerative disorders: parvocellular vs magnocellular degeneration in optical coherence tomography studies. Frontiers in neurology, 8:710, 2017a.

Chiara La Morgia, Fred N Ross-Cisneros, Alfredo A Sadun, and Valerio Carelli. Retinal ganglion cells and circadian rhythms in alzheimer’s disease, parkinson’s disease, and beyond. Frontiers in neurology, 8:162, 2017b.

Lidawani Lambuk, Mohd Aizuddin Mohd Lazaldin, Suhana Ahmad, Igor Iezhitsa, Renu Agarwal, Vuk Uskoković, and Rohimah Mohamud. Brain-derived neurotrophic factor-mediated neuroprotection in glaucoma: a review of current state of the art. Frontiers in Pharmacology, 13:875662, 2022.

Margaret Livingstone and David Hubel. Segregation of form, color, movement, and depth: anatomy, physiology, and perception. Science, 240(4853):740–749, 1988.

Niru Maheswaranathan, David B Kastner, Stephen A Baccus, and Surya Ganguli. Inferring hidden structure in multilayered neural circuits. PLoS computational biology, 14(8):e1006291, 2018.

Niru Maheswaranathan, Lane T McIntosh, Hidenori Tanaka, Satchel Grant, David B Kastner, Joshua B Melander, Aran Nayebi, Luke E Brezovec, Julia H Wang, Surya Ganguli, et al. Interpreting the retinal neural code for natural scenes: From computations to neurons. Neuron, 111(17):2742–2755, 2023.

Rania A Masri, Kumiko A Percival, Amane Koizumi, Paul R Martin, and Ulrike Grünert. Survey of retinal ganglion cell morphology in marmoset. Journal of Comparative Neurology, 527(1):236–258, 2019.

Lane McIntosh, Niru Maheswaranathan, Aran Nayebi, Surya Ganguli, and Stephen Baccus. Deep learning models of the retinal response to natural scenes. Advances in neural information processing systems, 29, 2016.

Markus Meister. A standard model for the vertebrate retina. Vision Research, 249:108895, 2026.

JO Mills, A Jalil, and PE Stanga. Electronic retinal implants and artificial vision: journey and present. Eye, 31(10):1383–1398, 2017.

Thomas W Nugent, Drew D Carter, Samir Uprety, Prakash Adhikari, Beatrix Feigl, and Andrew J Zele. Protocol for isolation of melanopsin and rhodopsin in the human eye using silent substitution. STAR protocols, 4(1), 2023.

Stefano Panzeri, Christopher D Harvey, Eugenio Piasini, Peter E Latham, and Tommaso Fellin. Cracking the neural code for sensory perception by combining statistics, intervention, and behavior. Neuron, 93(3):491–507, 2017.

Nikhil Parthasarathy, Eleanor Batty, William Falcon, Thomas Rutten, Mohit Rajpal, EJ Chichilnisky, and Liam Paninski. Neural networks for efficient bayesian decoding of natural images from retinal neurons. Advances in Neural Information Processing Systems, 30, 2017.

Yi-Rong Peng, Karthik Shekhar, Wenjun Yan, Dustin Herrmann, Anna Sappington, Gregory S Bryman, Tavé van Zyl, Michael Tri H Do, Aviv Regev, and Joshua R Sanes. Molecular classification and comparative taxonomics of foveal and peripheral cells in primate retina. Cell, 176(5):1222–1237, 2019.

Dumitru Petrusca, Matthew I Grivich, Alexander Sher, Greg D Field, Jeffrey L Gauthier, Martin Greschner, Jonathon Shlens, EJ Chichilnisky, and Alan M Litke. Identification and characterization of a y-like primate retinal ganglion cell type. Journal of Neuroscience, 27(41):11019–11027, 2007.

AJ Phillips, Amrith Lotlikar, Alexandra Kling, Madeline Hays, Praful Vasireddy, Bella Hofflich, Aviv Sharon, Michael A Sommeling, Jeff Brown, Claire Baum, et al. Electrical stimulation precisely reproduces naturalistic spiking activity in complete and intermixed neural populations in the primate retina. bioRxiv, pages 2026–05, 2026.

Pawel Pierzchlewicz, Konstantin Willeke, Arne Nix, Pavithra Elumalai, Kelli Restivo, Tori Shinn, Cate Nealley, Gabrielle Rodriguez, Saumil Patel, Katrin Franke, et al. Energy guided diffusion for generating neurally exciting images. Advances in Neural Information Processing Systems, 36:32574–32601, 2023.

Joel Pokorny and Vivianne C Smith. Psychophysical signatures associated with magnocellular and parvocellular pathway contrast gain. Journal of the Optical Society of America A, 14(9):2477–2486, 1997.

Carlos R Ponce, Will Xiao, Peter F Schade, Till S Hartmann, Gabriel Kreiman, and Margaret S Livingstone. Evolving images for visual neurons using a deep generative network reveals coding principles and neuronal preferences. Cell, 177(4):999–1009, 2019.

Jacob S Prince, Binxu Wang, Thomas Fel, Akshay V Jagadeesh, Parisa A Vaziri, George A Alvarez, Margaret S Livingstone, and Talia Konkle. Parametric neural control differentiates top neural network models of primate visual cortex. bioRxiv, pages 2026–08, 2026.

Harry A Quigley. Neuronal death in glaucoma. Progress in retinal and eye research, 18(1):39–57, 1999.

Suva Roy, Na Young Jun, Emily L Davis, John Pearson, and Greg D Field. Inter-mosaic coordination of retinal receptive fields. Nature, 592(7854):409–413, 2021.

Gregory W Schwartz, Haruhisa Okawa, Felice A Dunn, Josh L Morgan, Daniel Kerschensteiner, Rachel O Wong, and Fred Rieke. The spatial structure of a nonlinear receptive field. Nature neuroscience, 15(11):1572–1580, 2012.

Nishal P Shah, Nora Brackbill, Ryan Samarakoon, Colleen Rhoades, Alexandra Kling, Alexander Sher, Alan Litke, Yoram Singer, Jonathon Shlens, and EJ Chichilnisky. Individual variability of neural computations in the primate retina. Neuron, 110(4):698–708, 2022.

Scott M Sternson and Bryan L Roth. Chemogenetic tools to interrogate brain functions. Annual review of neuroscience, 37(1):387–407, 2014.

Ian Van Der Linde, Umesh Rajashekar, Alan C Bovik, and Lawrence K Cormack. Doves: a database of visual eye movements. Spatial vision, 22(2):161, 2009.

J Hans Van Hateren and Arjen van der Schaaf. Independent component filters of natural images compared with simple cells in primary visual cortex. Proceedings of the Royal Society B: Biological Sciences, 265(1394):359, 1998.

Edgar Y Walker, Fabian H Sinz, Erick Cobos, Taliah Muhammad, Emmanouil Froudarakis, Paul G Fahey, Alexander S Ecker, Jacob Reimer, Xaq Pitkow, and Andreas S Tolias. Inception loops discover what excites neurons most using deep predictive models. Nature neuroscience, 22(12):2060–2065, 2019.

H Wässle, Brian Blundell Boycott, and R-B Illing. Morphology and mosaic of on-and off-beta cells in the cat retina and some functional considerations. Proceedings of the Royal Society of London. Series B. Biological Sciences, 212(1187):177–195, 1981.

Heinz Wassle and BRIAN B Boycott. Functional architecture of the mammalian retina. Physiological reviews, 71(2):447–480, 1991.

Anqi Wu, Il Memming Park, and Jonathan W Pillow. Convolutional spike-triggered covariance analysis for neural subunit models. Advances in neural information processing systems, 28, 2015.

Elizabeth S Yamada, Andrea S Bordt, and David W Marshak. Wide-field ganglion cells in macaque retinas. Visual neuroscience, 22(4):383–393, 2005.

Daniel LK Yamins, Ha Hong, Charles F Cadieu, Ethan A Solomon, Darren Seibert, and James J DiCarlo. Performance-optimized hierarchical models predict neural responses in higher visual cortex. Proceedings of the national academy of sciences, 111(23):8619–8624, 2014.

