## Supplementary Data for "Designing stimuli for non-invasive, cell-type-specific manipulation of retinal ganglion cell activity"

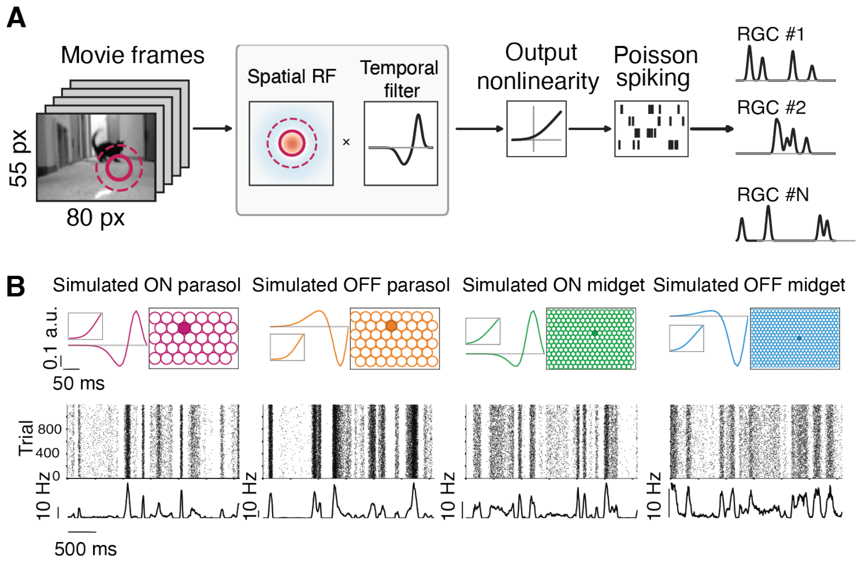


**Figure S1. Parameters of simulated RGC types.** **(A)** Schematic of the LNP model, as in Fig. 2A. **(B)** Properties of four simulated RGC types. For each type: temporal filter (top left), RF mosaic (top right), output nonlinearity (inset), and spike rasters across repeated trials with the corresponding trial-averaged firing rate (bottom).

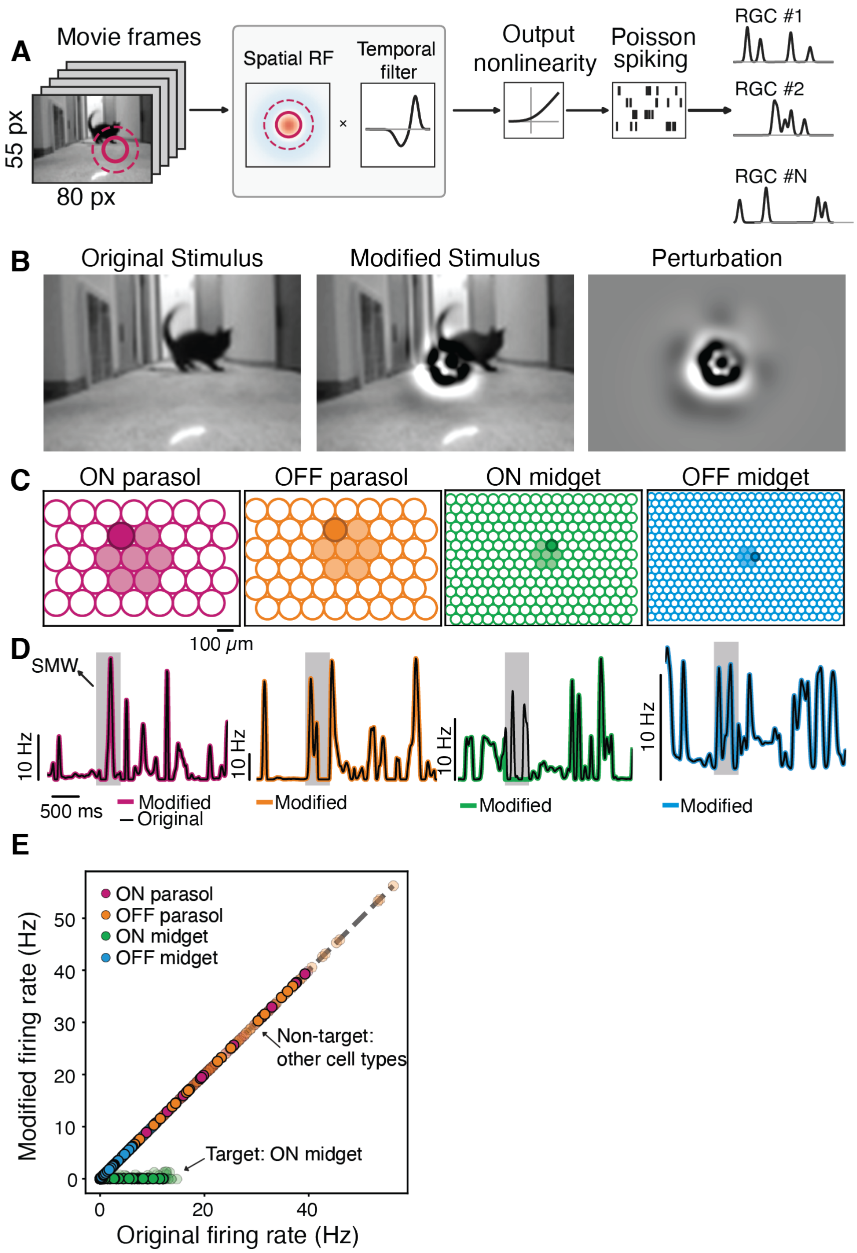


**Figure S2. Selective suppression of simulated ON midget responses using a Linear-Nonlinear-Poisson model. (A)** Schematic of the LNP model, as in Fig. 2A. **(B)** Example frame of the original movie (left), the corresponding frame of the modified movie that suppresses ON midget cells (middle), and the difference between the modified and original frames (perturbation; right). **(C)** RF mosaics of the simulated RGC types drawn at 1-$\sigma$. Filled RFs indicate the cells included in the manipulation; darker cells are shown in **D**. **(D)** Firing rates for example cells in response to the original (black) and modified (colored) movies; the stimulus manipulation window (SMW) is shaded gray. The ON midget response is reduced during the SMW, the non-target responses are unmodified. **(E)** Firing rate of every cell at each time bin (each point) within the SMW for the original vs modified movies. ON midget cells are suppressed to near-zero rate, the three non-target types are unmodulated. Opaque points are the four cells shown in **D**.

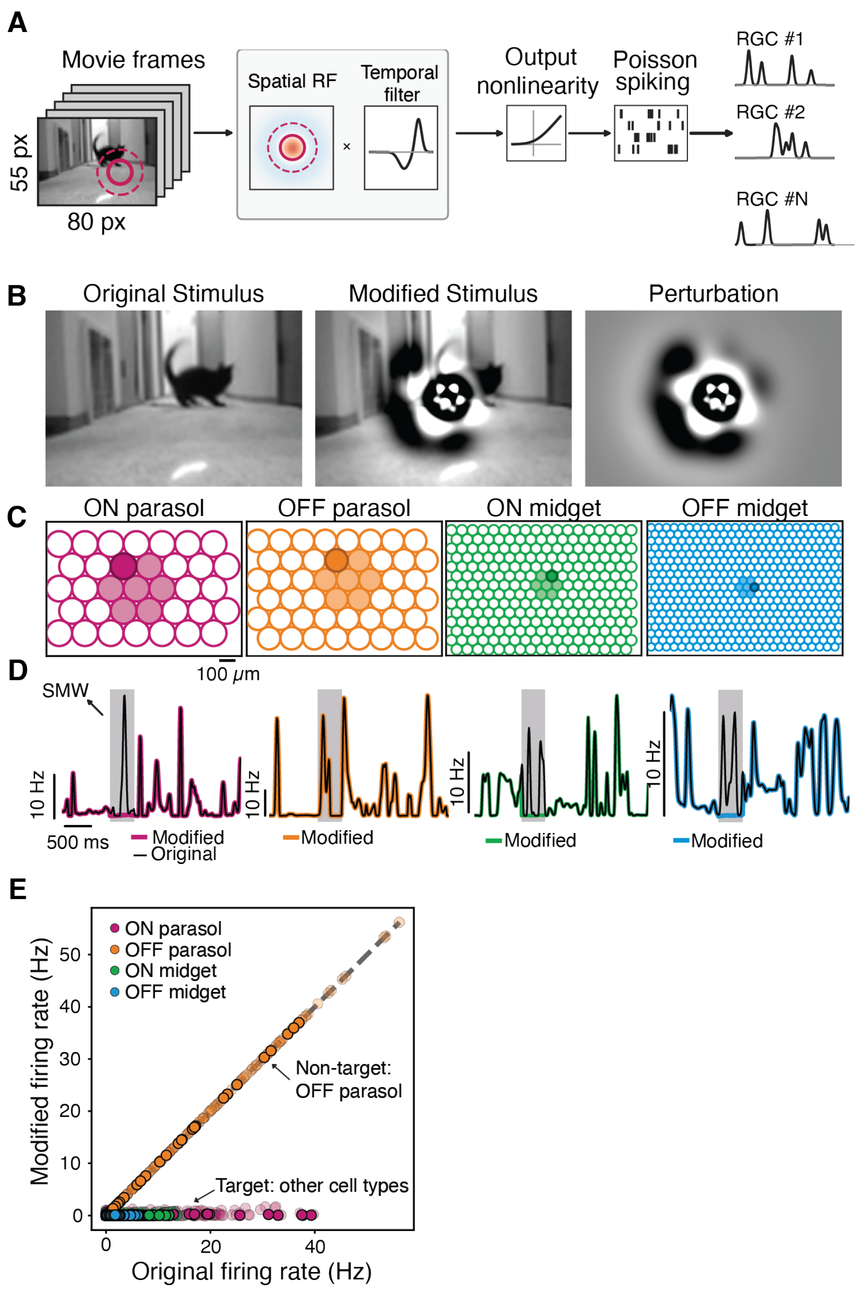


**Figure S3. Simultaneous suppression of multiple simulated RGC type responses using a Linear-Nonlinear-Poisson model.** **(A)** Schematic of the LNP model, as in Fig. 2A. **(B)** Example frame of the original movie (left), the corresponding frame of the modified movie that suppresses ON parasol, ON and OFF midget cells (middle), and the difference between the modified and original frames (perturbation; right). **(C)** RF mosaics of the simulated RGC types drawn at 1-$\sigma$. Filled RFs indicate the cells included in the manipulation; darker cells are shown in **D**. **(D)** Firing rates for example cells in response to the original (black) and modified (colored) movies; the stimulus manipulation window (SMW) is shaded gray. The ON parasol, ON and OFF midget cells responses are reduced during the SMW, OFF parasol responses are unmodified. **(E)** Firing rate of every cell at each time bin (each point) within the SMW for the original vs modified movies. ON parasol, ON and OFF midget cells are suppressed to near-zero rates, OFF parasol responses are unmodified. Opaque points are the four cells shown in **D**.

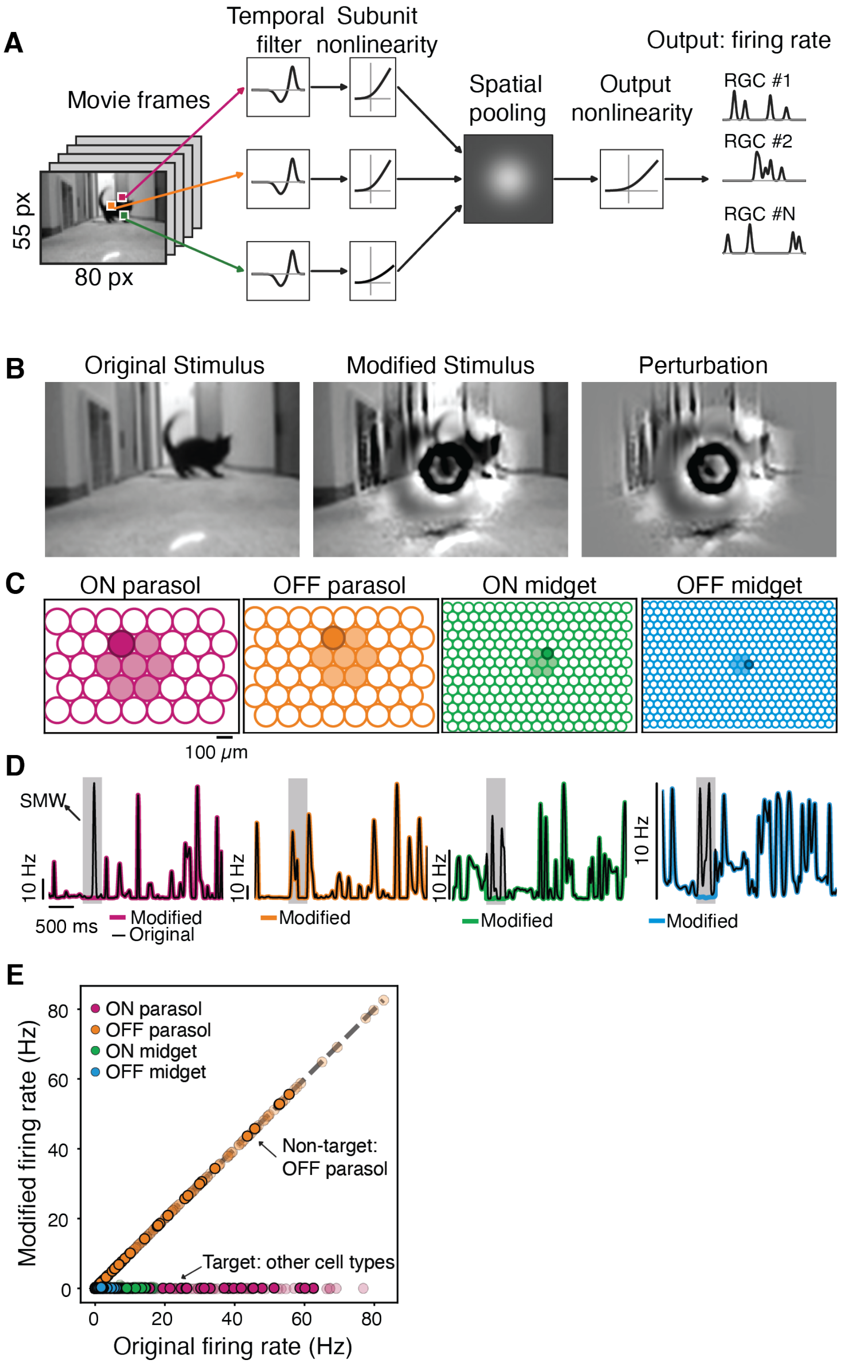


**Figure S4. Simultaneous suppression of multiple simulated RGC type responses using a Linear-Nonlinear-Linear-Nonlinear model.** **(A)** Schematic of the LN-LN model, as in Fig. 3A. **(B)** Example frame of the original movie (left), the corresponding frame of the modified movie that suppresses ON parasol, ON and OFF midget cells (middle), and the difference between the modified and original frames (perturbation; right). **(C)** RF mosaics of the simulated RGC types drawn at 1-$\sigma$. Filled RFs indicate the cells included in the manipulation; darker cells are shown in **D**. **(D)** Firing rates for example cells in response to the original (black) and modified (colored) movies; the stimulus manipulation window (SMW) is shaded gray. The ON parasol, ON and OFF midget cells responses are reduced during the SMW, OFF parasol responses are unmodified. **(E)** Firing rate of every cell at each time bin (each point) within the SMW for the original vs modified movies. ON parasol, ON and OFF midget cells are suppressed to near-zero rates, OFF parasol responses are unmodified. Opaque points are the four cells shown in **D**.

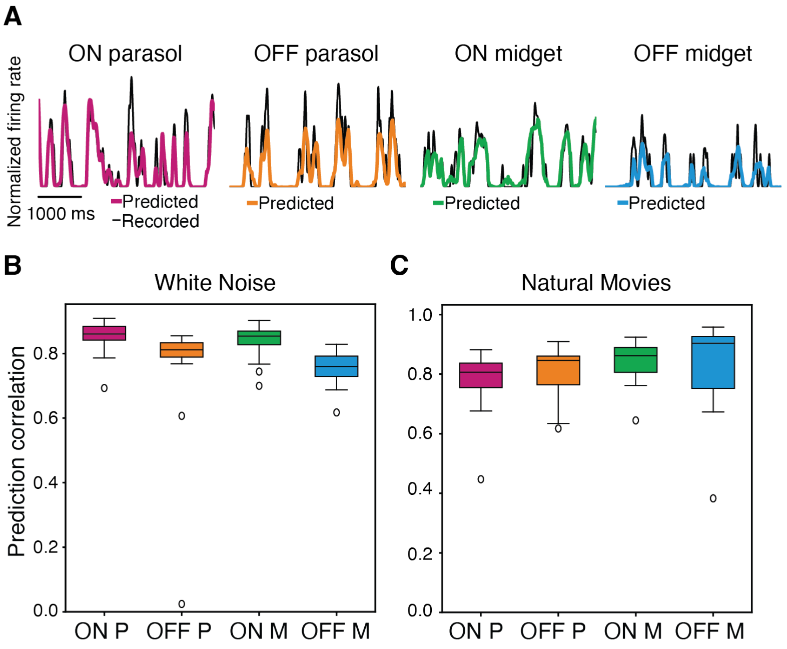


**Figure S5. Predictive performance of the CNN model.** **(A)** Recorded (black) and predicted (colored) firing rates for one example RGC of each type in response to a held-out movie segment. **(B)** Pearson correlation between predicted and recorded firing rates for each RGC type for the model trained and tested on white-noise movies (ON parasol, n = 16; OFF parasol, n = 13; ON midget, n = 40; OFF midget, n = 15). Box plots show the median and interquartile range; whiskers extend to the most extreme values within 1.5 times the interquartile range and open circles denote outliers. **(C)** Same as **B**, for the model trained on white noise and subsequently fine-tuned on natural movies, evaluated on held-out natural-movie segments.

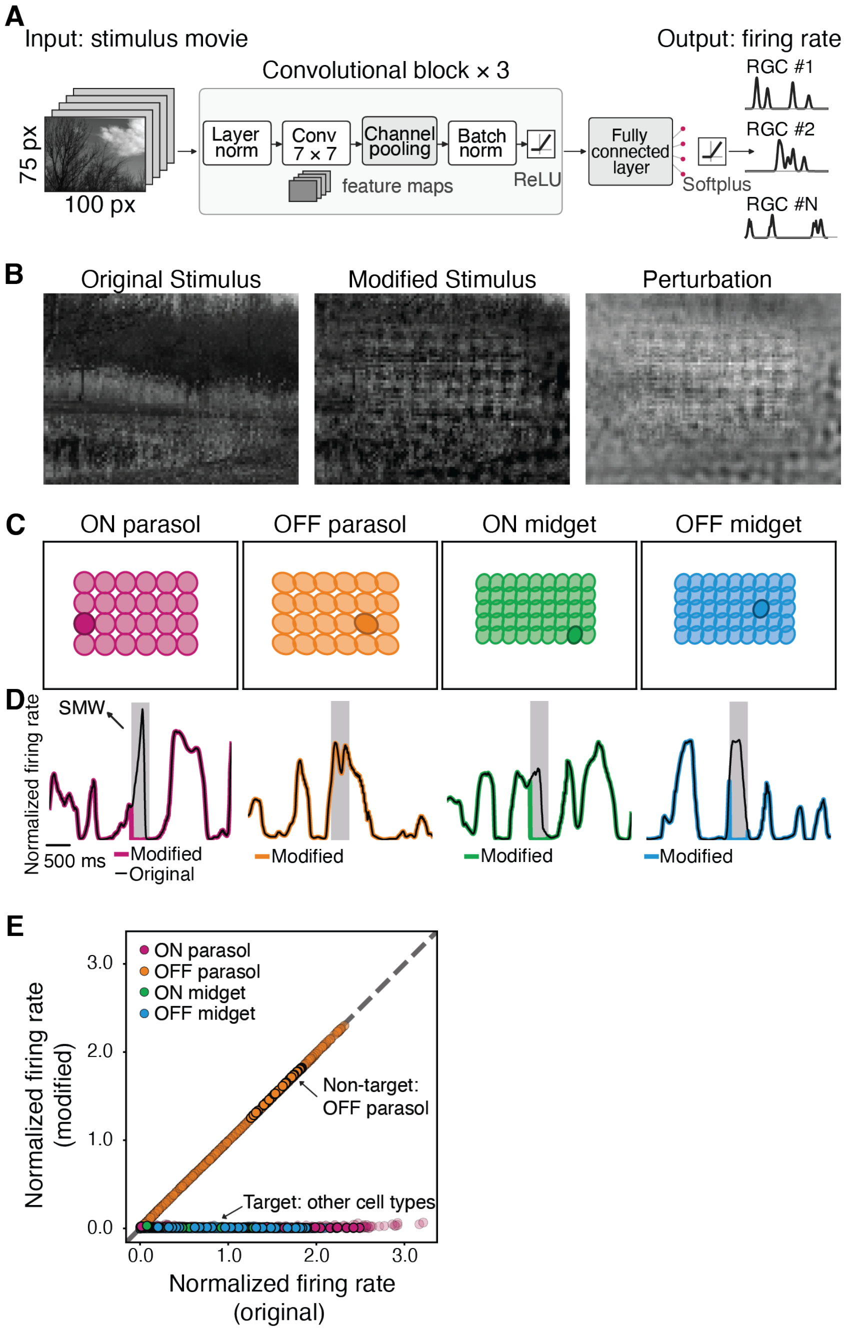


**Figure S6. Simultaneous suppression of multiple simulated RGC type responses using a CNN model.** **(A)** Schematic of the CNN model, as in Fig.4A. **(B)** Example frame of the original movie (left), the corresponding frame of the modified movie that suppresses ON parasol, ON and OFF midget cells (middle), and the difference between the modified and original frames (perturbation; right). **(C)** RF mosaics of the simulated types drawn at 1-$\sigma$. Filled RFs indicate the cells included in the manipulation; darker cells are shown in **D**. **(D)** Firing rates for example cells in response to the original (black) and modified (colored) movies; the stimulus manipulation window (SMW) is shaded gray. The ON parasol, ON and OFF midget cells responses are reduced during the SMW, OFF parasol responses are unmodified. **(E)** Firing rate of every cell at each time bin (each point) within the SMW for the original vs modified movies. ON parasol, ON and OFF midget cells are suppressed to near-zero rates, OFF parasol responses are unmodified. Opaque points are the four cells shown in **D**.

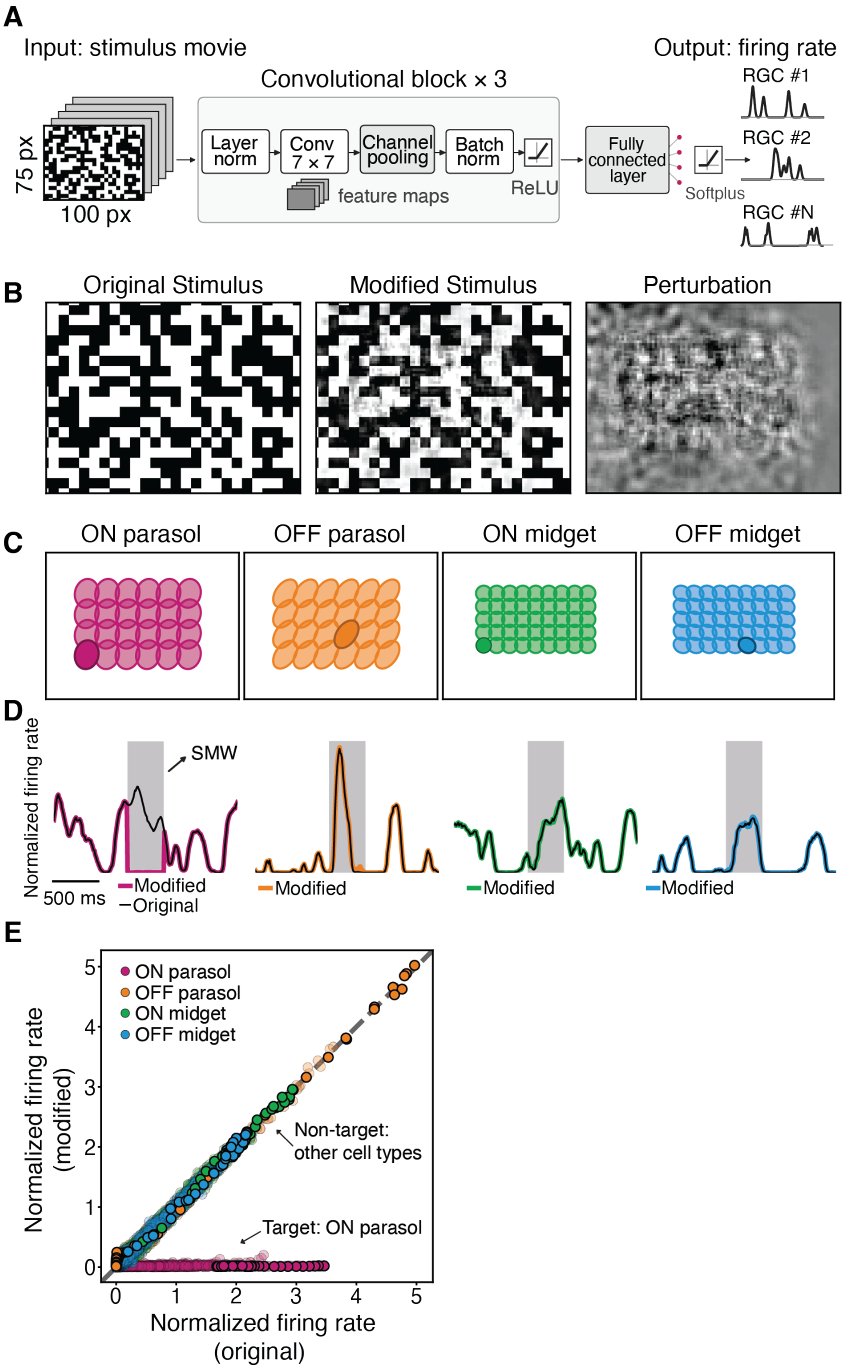


**Figure S7. Selective suppression of simulated RGC responses to a white noise checkerboard movie using a CNN model.** **(A)** Schematic of the CNN model, as in Fig.4A. **(B)** Example frame of the original white noise checkerboard movie (left), the corresponding frame of the modified movie that suppresses ON parasol cells (middle), and the difference (perturbation, right). **(C)** RF mosaics of the simulated types drawn at 1-$\sigma$. Filled RFs indicate the cells included in the manipulation; darker cells are shown in **D**. **(D)** Firing rates normalized by the median firing rate for example cells in response to the original (black) and modified (colored) movies; the stimulus manipulation window (SMW) is shaded gray. The ON parasol response is reduced during the SMW; the non-target responses are unmodified. **(E)** Firing rate of every cell at each time bin (each point) within the SMW for the original vs modified movies. ON parasol cells are suppressed to near-zero rates; the three non-target types are unmodulated. Opaque points are the four cells shown in **D**.

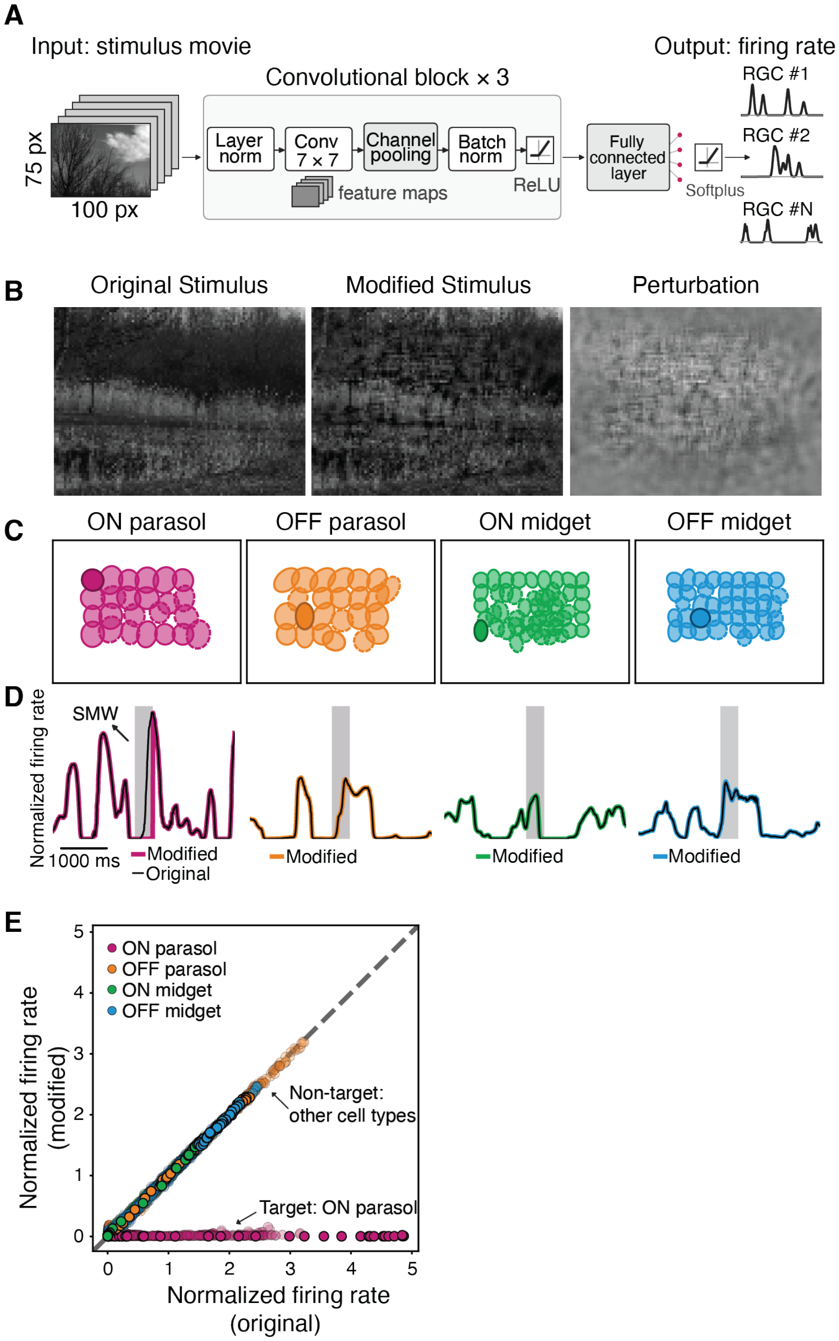


**Figure S8. Selective suppression of simulated responses of real RGCs in a heterogeneous mosaic using a CNN model. (A)** Schematic of the CNN model, as in Fig.4A. **(B)** Example frame of the original natural movie (left), the corresponding frame of the modified movie that suppresses ON parasol cells (middle), and the difference (perturbation, right). **(C)** RF mosaics of the simulated types drawn at 1-$\sigma$. Filled RFs indicate the cells included in the manipulation; darker cells are shown in **D**. Recorded RGCs are shown with dashed outlines and synthetic RGCs with solid outlines. **(D)** Firing rates normalized by the median firing rate for example cells in response to the original (black) and modified (colored) movies; the stimulus manipulation window (SMW) is shaded gray. The ON parasol response is reduced during the SMW; the non-target responses are unmodified. **(E)** Firing rate of every cell at each time bin (each point) within the SMW for the original vs modified movies. ON parasol cells are suppressed to near-zero rates; the three non-target types are unmodulated. Opaque points are the four cells shown in **D**.

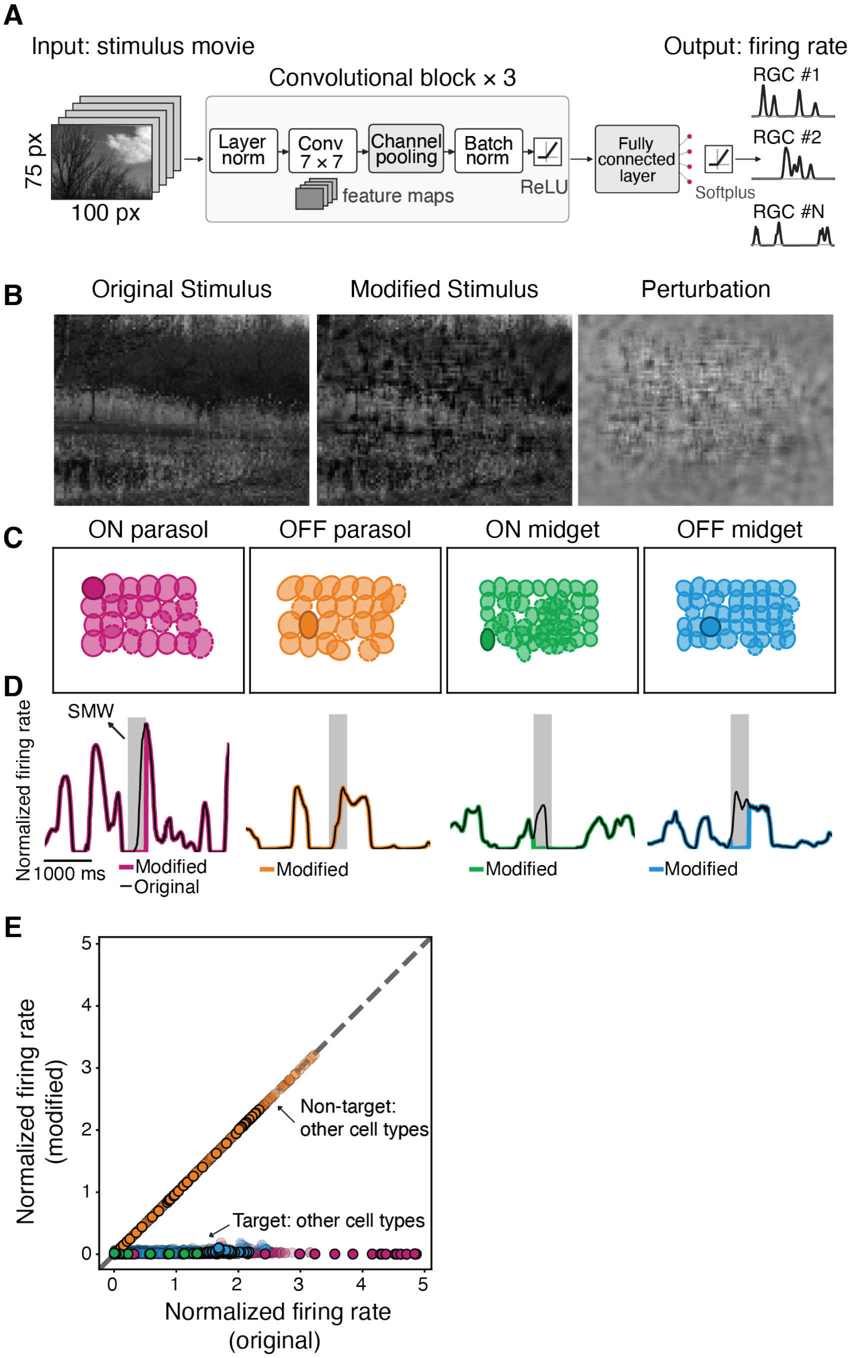


**Figure S9. Simultaneous suppression of multiple simulated responses of real RGCs in a heterogeneous mosaic using a CNN model.** **(A)** Schematic of the CNN model, as in Fig.4A. **(B)** Example frame of the original movie (left), the corresponding frame of the modified movie that suppresses ON parasol, ON and OFF midget cells (middle), and the difference between the modified and original frames (perturbation; right). **(C)** RF mosaics of the simulated types drawn at 1-$\sigma$. Filled RFs indicate the cells included in the manipulation; darker cells are shown in **D**.Recorded RGCs are shown with dashed outlines and synthetic RGCs with solid outlines. **(D)** Firing rates for example cells in response to the original (black) and modified (colored) movies; the stimulus manipulation window (SMW) is shaded gray. The ON parasol, ON and OFF midget cells responses are reduced during the SMW, OFF parasol responses are unmodified. **(E)** Firing rate of every cell at each time bin (each point) within the SMW for the original vs modified movies. ON parasol, ON and OFF midget cells are suppressed to near-zero rates, OFF parasol responses are unmodified. Opaque points are the four cells shown in **D**.

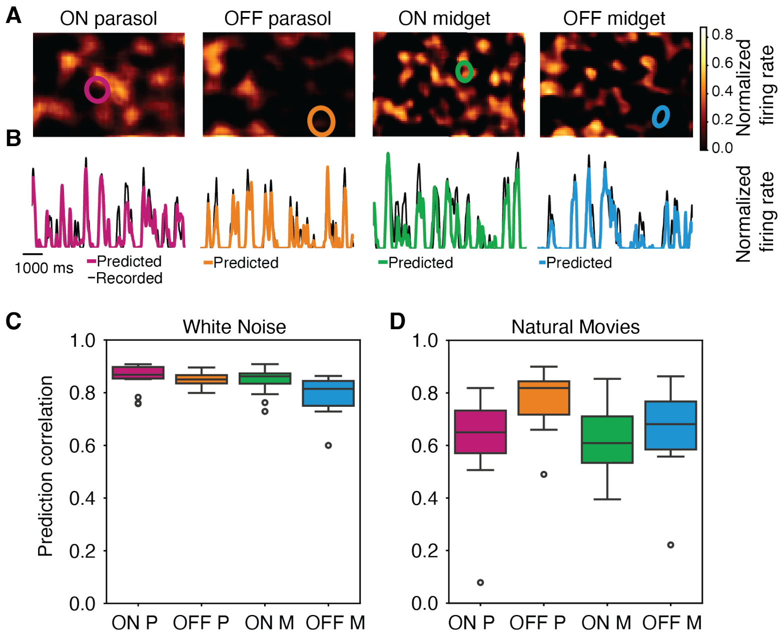


**Figure S10. Predictive performance of the map-based CNN model. (A)** Predicted firing-rate maps for ON parasol, OFF parasol, ON midget and OFF midget cells at a single time point. The colored circle in each map marks the RF center of one recorded RGC of that type; the predicted response at that location is compared with the recorded response of that cell in **B**. **(B)** Recorded (black) and predicted (colored) firing rates over time at the locations marked in **A** for a held-out white-noise segment. **(C)** Pearson correlation between predicted and recorded firing rates for the recorded cells of each type for the model trained and tested on white-noise movies. Box plots show the median and interquartile range; whiskers extend to the most extreme values within 1.5 times the interquartile range and open circles denote outliers. **(D)** Same as **C**, for the model trained on white noise and evaluated on natural movies.

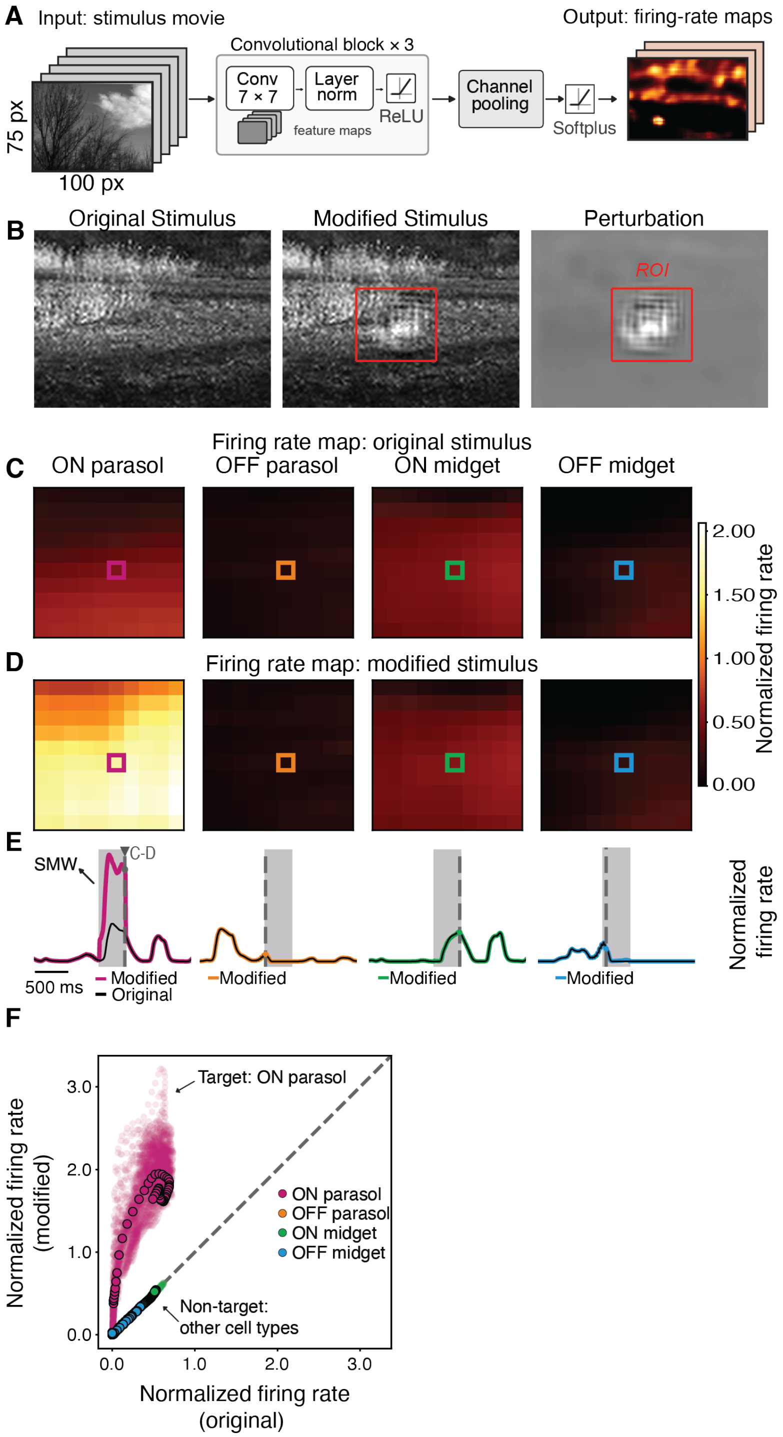


**Figure S11. Selective enhancement of RGC responses with translation invariance using a CNN and firing rate maps. (A)** Schematic of the map-based CNN as in Fig. 5A. **(B)** Example frame of the original natural movie (left), the corresponding frame of the modified movie that enhances ON parasol cells (middle), and the difference (perturbation, right). The outlined region denotes the region of interest (ROI) analyzed in **C-F**. **(C)** Firing-rate maps within the ROIs of **B** for each RGC type in response to the original movie, at the time indicated by the dashed vertical line in **E**. **(D)** Same as **C**, for the modified movie. The ON parasol firing rate map is enhanced within the ROI, whereas the other three non-target RGC types are unaffected. **(E)** Predicted firing rate over time at the locations marked by the colored squares in **C-D**, normalized by the maximum firing rate, in response to the original (black) and modified (colored) movies. Gray shading marks the SMW; the dashed vertical line marks the time point of the maps shown in **C-D**. **(F)** Scatter plot compares firing rate during the SMW at every location within the ROI, in response to the original and the modified movie. ON parasol responses are enhanced at all locations; the non-target RGC responses are unchanged. Opaque points are the four cells shown in **E**.

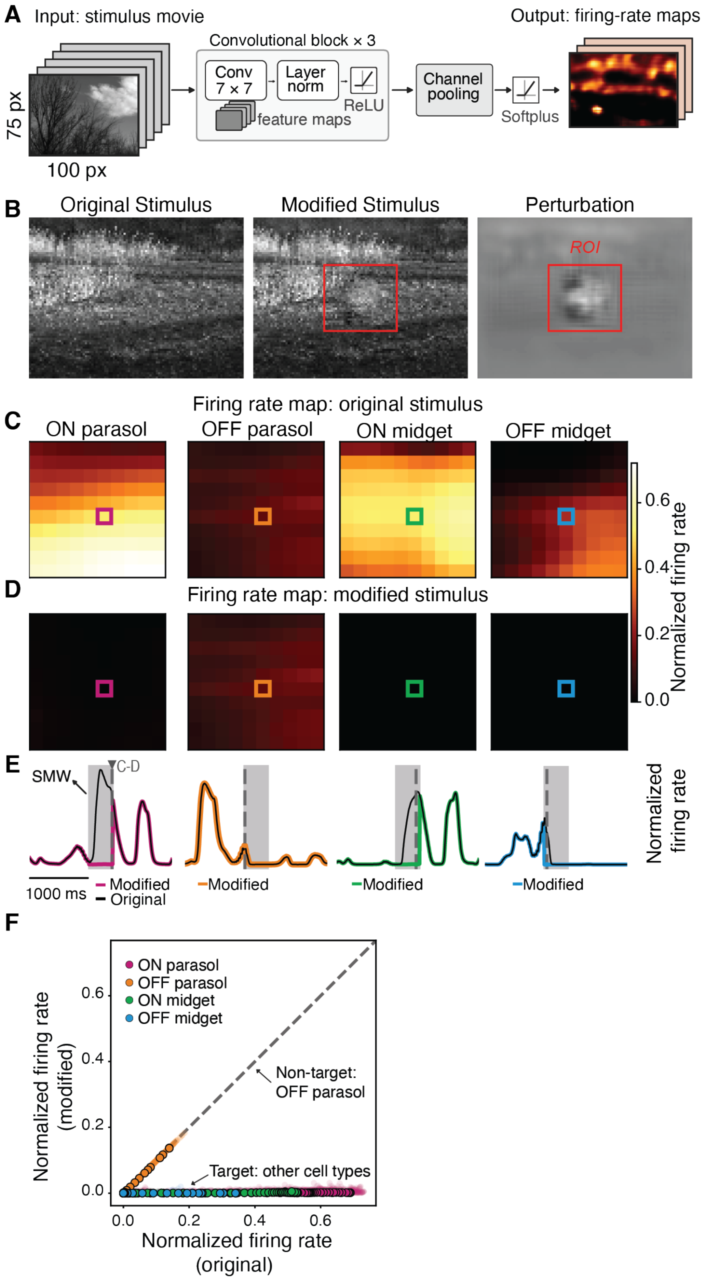


**Figure S12. Simultaneous suppression of multiple RGC type responses with translation invariance using a CNN and firing rate maps.** **(A)** Schematic of the map-based CNN as in Fig. 5A. **(B)** Example frame of the original natural movie (left), the corresponding frame of the modified movie that suppresses ON parasol, ON and OFF midget cells (middle), and the difference (perturbation, right). The outlined region denotes the region of interest (ROI) analyzed in **C-F**. **(C)** Firing-rate maps within the ROIs of **B** for each RGC type in response to the original movie, at the time indicated by the dashed vertical line in **E**. **(D)** Same as **C**, for the modified movie. The ON parasol, ON and OFF midget firing rate maps are driven to near-zero within the ROI, whereas the non-target RGC type (OFF parasol) is unaffected. **(E)** Predicted firing rate over time at the locations marked by the colored squares in **C-D**, normalized by the maximum firing rate, in response to the original (black) and modified (colored) movies. Gray shading marks the SMW; the dashed vertical line marks the time point of the maps shown in **C-D**. **(F)** Scatter plot compares firing rate during the SMW at every location within the ROI, in response to the original and the modified movie. The ON parasol, ON and OFF midget cells are suppressed to near-zero rates, OFF parasol responses are unmodified. Opaque points are the four cells shown in **E**.

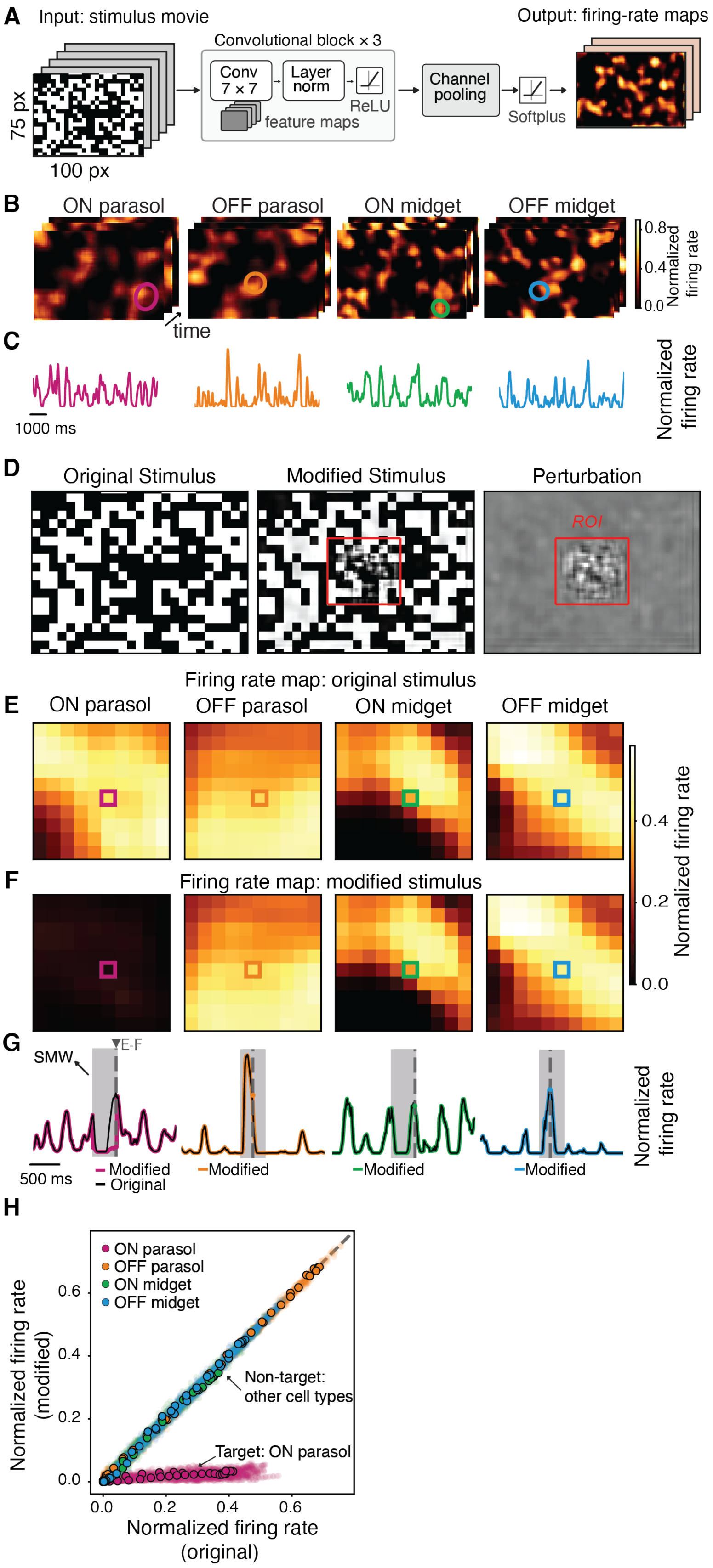


**Figure S13. Selective RGC manipulation with translation invariance using a CNN and firing rate maps with a white noise checkerboard movie.** **(A)** Schematic of the map-based CNN as in Fig. 5A. **(B)** Example firing-rate maps for the four RGC types at successive time points. The colored circles mark RF centers over which a firing rate can be integrated. **(C)** Predicted firing rate over time at the RF locations in **B**. **(B)** Example frame of the original white noise checkerboard movie (left), the corresponding frame of the modified movie that suppresses ON parasol cells (middle), and the difference (perturbation, right). The outlined region denotes the region of interest (ROI) analyzed in **E-H**. **(E)** Firing-rate maps within the ROIs of **D** for each RGC type in response to the original movie, at the time indicated by the dashed vertical line in **G**. **(F)** Same as **E**, for the modified movie. The ON parasol firing rate map is driven to near-zero within the ROI, whereas the other three non-target RGC types are unaffected. **(G)** Predicted firing rate over time at the locations marked by the colored squares in **E-F**, normalized by the maximum firing rate, in response to the original (black) and modified (colored) movies. Gray shading marks the SMW; the dashed vertical line marks the time point of the maps shown in **E-F**. **(H)** Scatter plot compares firing rate during the SMW at every location within the ROI, in response to the original and the modified movie. ON parasol responses are suppressed at all locations; the non-target RGC responses are unchanged. Opaque points are the four cells shown in **G**.

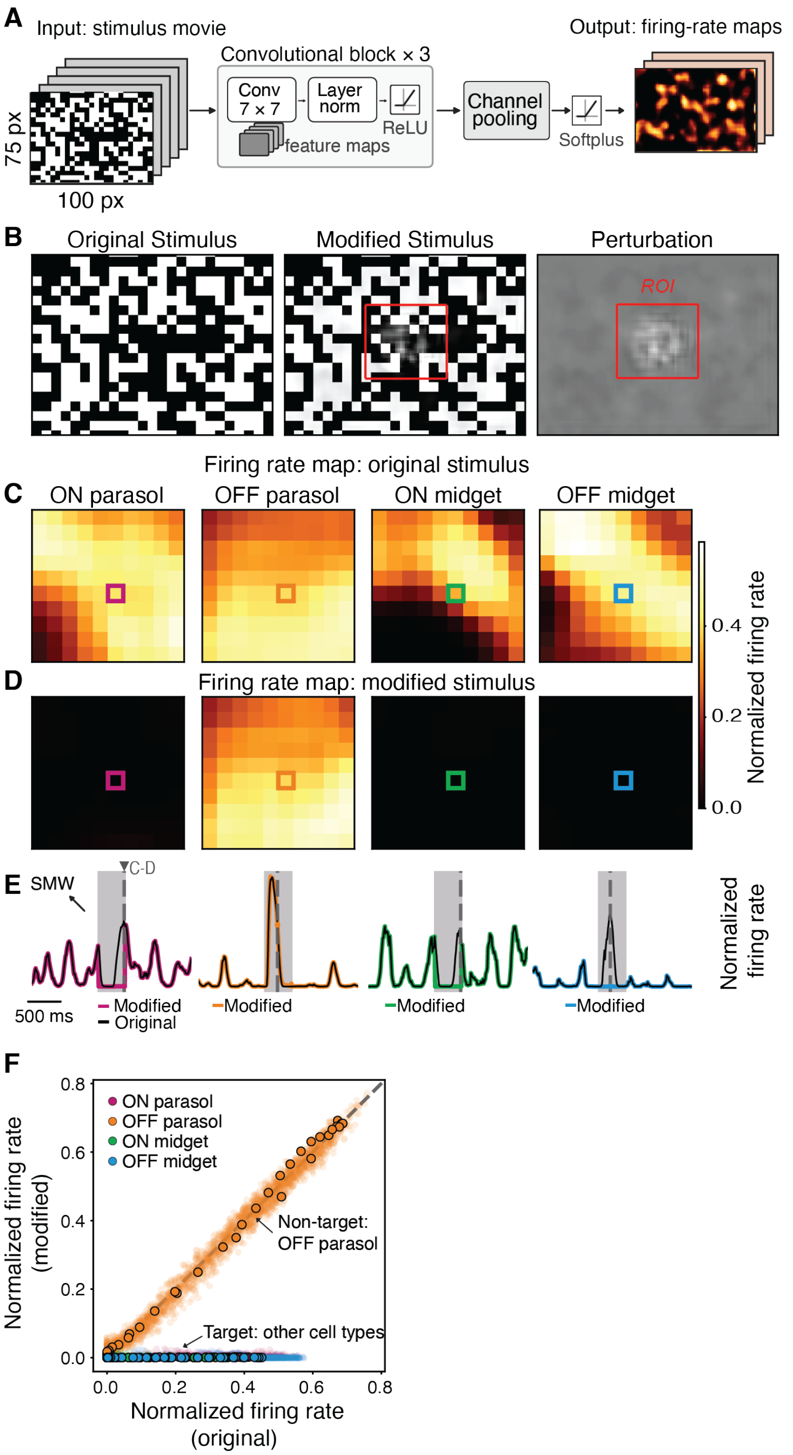


**Figure S14. Simultaneous suppression of multiple RGC type responses with translation invariance using a CNN and firing rate maps with a white noise checkerboard movie.** **(A)** Schematic of the map-based CNN as in Fig. 5A. **(B)** Example frame of the original white noise checkerboard movie (left), the corresponding frame of the modified movie that suppresses ON parasol, ON and OFF midget cells (middle), and the difference (perturbation, right). The outlined region denotes the region of interest (ROI) analyzed in **C-F**. **(C)** Firing-rate maps within the ROIs of **B** for each RGC type in response to the original movie, at the time indicated by the dashed vertical line in **E**. **(D)** Same as **C**, for the modified movie. The ON parasol, ON and OFF midget firing rate maps are driven to near-zero within the ROI, whereas the non-target RGC type (OFF parasol) is unaffected. **(E)** Predicted firing rate over time at the locations marked by the colored squares in **C-D**, normalized by the maximum firing rate, in response to the original (black) and modified (colored) movies. Gray shading marks the SMW; the dashed vertical line marks the time point of the maps shown in **C-D**. **(F)** Scatter plot compares firing rate during the SMW at every location within the ROI, in response to the original and the modified movie. The ON parasol, ON and OFF midget cells are suppressed to near-zero rates, OFF parasol responses are unmodified. Opaque points are the four cells shown in **E**.

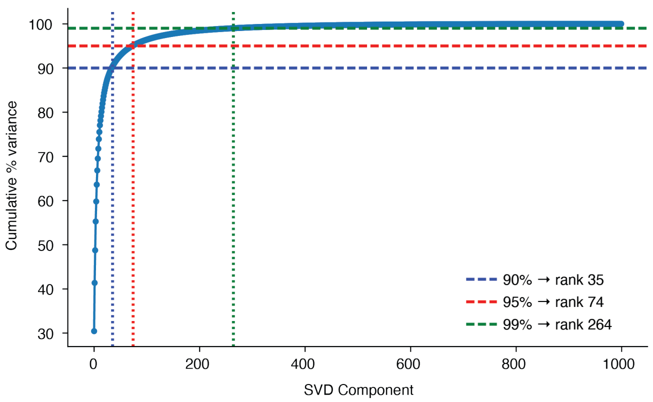


**Figure S15. Low-rank approximation of the non-target sensitivity subspace.** Cumulative percentage of variance in the non-target response gradients explained as a function of the number of retained singular value decomposition (SVD) components. Dashed horizontal and vertical lines mark the ranks required to capture 90% (blue, 35 components), 95% (red, 74 components) and 99% (green, 264 components) of the variance.
